# The lincRNA *Pantr1* is a FOXG1 target gene conferring site-specific chromatin binding of FOXG1

**DOI:** 10.1101/2024.08.29.610239

**Authors:** Fabian Gather, Tudor Rauleac, Ipek Akol, Ganeshkumar Arumugam, Camila L. Fullio, Teresa Müller, Dimitrios Kleidonas, Andre Fischer, Andreas Vlachos, Rolf Backofen, Tanja Vogel

## Abstract

Derailed gene expression programs within the developing nervous system, encompassing both transcriptional and posttranscriptional processes, can cause diverse neurodevelopmental diseases (NDD). The NDD FOXG1-syndrome lacks full understanding of the mechanistic role of its eponymous gene product. While it is known that FOXG1 acts in part at the chromatin by binding to regulative regions, it is unclear what factors control its presence at specific sites. Long non-coding RNAs (lncRNAs) can mediate site-directed transcription factor binding, but their potential role in FOXG1-syndrome has not been described. Here, we show that FOXG1 localisation is regulated at selected loci through the lncRNA *Pantr1*.

We identified FOXG1 as an upstream transcriptional activator of *Pantr1* in human and mice. Further, we discovered that FOXG1 has the ability to associate with RNAs. Both, transcriptional regulation of *Pantr1* by FOXG1 and association of both partners, build up a regulative network that impacts the localisation of FOXG1 at selected genomic loci. Specifically, *Pantr1* facilitates cooperative presence of FOXG1/NEUROD1 at specific sites, and *Pantr1* reduction leads to redistribution of FOXG1 to comparably more generic binding sites. The rescue of impaired dendritic outgrowth upon FOXG1 reduction by simultaneous overexpression of *Pantr1* underlines the importance of the FOXG1/*Pantr1* regulative network.

**GRAPHICAL ABSTRACT:** 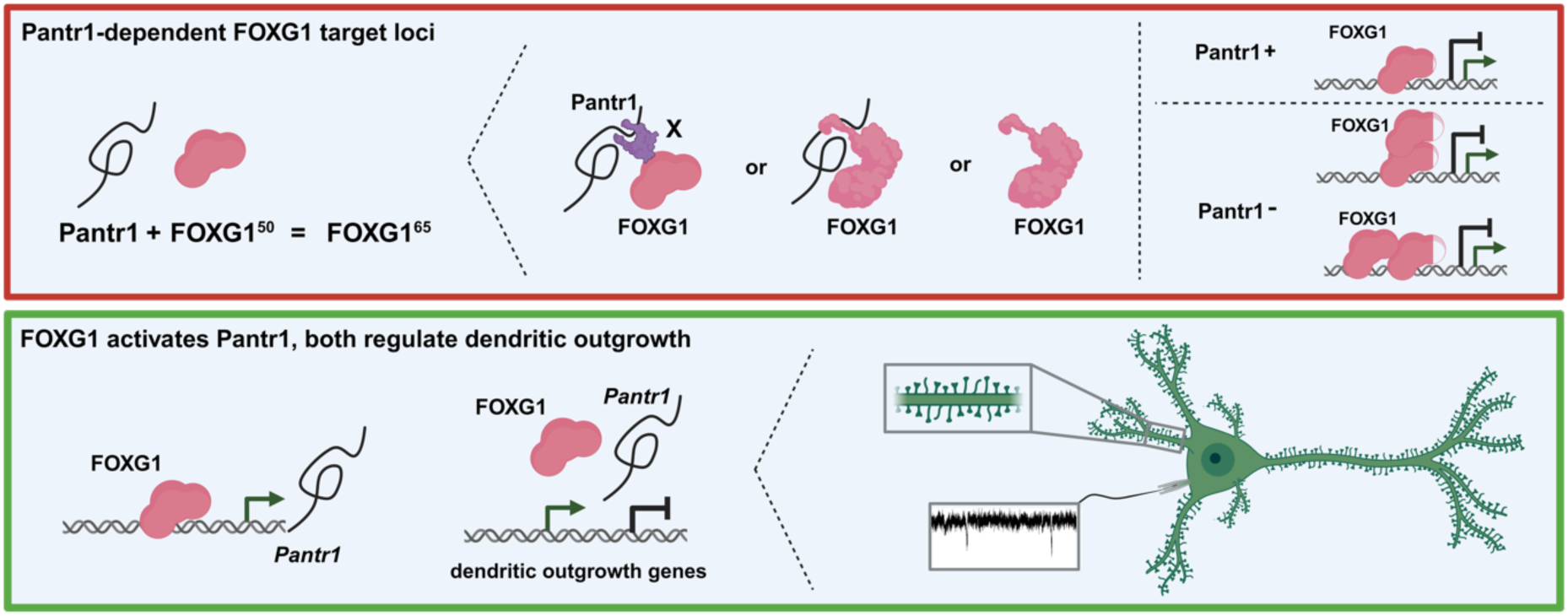

## INTRODUCTION

Forkhead box G1 (FOXG1)-syndrome (OMIM #613454) is a rare, congenital form in the Rett- and autism-spectrum disorders. Patients with loss-of-function *FOXG1* mutations present with a complex phenotypic spectrum (1), caused at least in part by impaired neuronal networks and functions. FOXG1’s link to autism is established both in humans (2) and mice (3). In mice, FOXG1 plays a central role in forebrain development as its complete absence results in anencephaly (4). It influences proliferation as well as differentiation of neural stem cells, and is involved in migration and integration of pyramidal neurons into the cortical plate (5-7). FOXG1 prevents neural stem cells from premature differentiation, thereby maintaining the number of progenitors available for the generation of specific neuronal subtypes. Accordingly, overexpression of FOXG1 increases the stem cell pool and delays neurogenesis (8). Moreover, FOXG1 expression inhibits differentiation of Cajal-Retzius-cells, which are overabundant in FOXG1-deficient forebrains (9), and FOXG1 deficiency impairs proper development of the ventral telencephalon through altered expression of ventralising signals (10).

FOXG1 also impacts the postnatal brain function. In the mouse postnatal hippocampus, for example, FOXG1 is involved in proper development of the dentate gyrus. Here, it maintains the balance between proliferation and differentiation of both neurons and glia cells (11), similar to that of cortical stem cells (8). Additionally, loss of FOXG1 results in increased cell death of postmitotic neurons in the hippocampus and cerebellum (11,12). Reduced levels of FOXG1 in the hippocampus further impact the animal’s behaviour. *Foxg1*-haploinsufficient mice are hyperactive, and they perform less well in contextual fear conditioning compared to wild-type controls (13). Recent studies show altered electrophysiological properties upon reduced or increased FOXG1 levels, dysbalanced neuronal functions of excitation and inhibition as important basis for impaired behaviour, reduced dendritic complexity and reduced spine densities, both in the cortex and hippocampus (3,14-17).

At the molecular level, the functions of FOXG1 are starting to gain recognition. FOXG1 prevents cell cycle exit of progenitor cells and promotes stem cell pool expansion by repressing the expression of Cyclin-dependent kinase 1a (*Cdkn1a)* (18,19). In this setting, FOXG1 sponges FOXO/SMAD complexes and prevents their binding to regulative sites, which normally activate *Cdkn1a* expression. Thereby, FOXG1 antagonises FOXO/SMAD-dependent neuronal differentiation of cortical progenitors (20). A growing number of studies reports on FOXG1 binding at the chromatin and its potential direct chromatin-related functions (6,7,21,22). Chromatin-related functions of FOXG1 impact gene expression programs that are implicated in neuronal differentiation, but also confer layer identity through its presence at a monomethylated histone H3 lysine 4 (H3K4me1) enhancer of the cortical layer 4 gene *Nr2f1*, which is kept in a repressed state. Removal of FOXG1 in projection neurons activates *Nr2f1* and a layer 4 neuron-specific transcriptional program (6). In addition, FOXG1 regulates callosal axon guidance and neuronal migration by acting within a FOXG1-ZBTB18 complex that directly binds and represses *Robo1, Slit3*, and *Reelin* genes (7).

While ChIP-seq data reveals FOXG1’s utilisation of diverse modalities to bind to chromatin and alter the epigenetic and transcriptional landscape (22), the mechanism by which FOXG1 discriminates its target chromatin regions remains unclear. Here, non-coding RNAs might confer sequence-specific recruitment of FOXG1. But despite the fact that specific miRNAs are involved in FOXG1-syndrome (23), it is not yet reported whether other non-coding RNAs, especially long (intergenic) non-coding RNAs (l(i)ncRNAs) contribute to this neurodevelopment disorder. This study unravels lncRNA, i.e. *Pantr1* contribution to FOXG1 syndrome, in both human and mouse model systems. We show that *Pantr1* is a direct target gene of FOXG1 and we unveil a novel mechanism mediating site-specific presence of FOXG1 at the chromatin level based on the association with *Pantr1*. Notably, the overexpression of *Pantr1* rescued dendritic outgrowth defects resulting from reduced FOXG1 expression. Our findings thus underscore the crucial role of the lincRNA *Pantr1* in FOXG1-mediated neuronal differentiation and function.

## MATERIAL AND METHODS

### Organoid culture

hiPSC cell lines were grown in mTESR1 medium (Stem Cell, Canada) on Matrigel (Corning, USA). Control hiPSCs (FiPS Core, University Medicine Freiburg, Germany) were generated from fibroblast of a healthy donor (24). FOXG1del hiPSCs were obtained from the Telethon, University of Siena (code 611/2013 #5) (17). hCO were generated according to an adapted version of the standard protocol (25), as described (26). Briefly, 35.000 cells per well of a v-shaped, non-coated 96 well plate were seeded, centrifuged and treated as described. The organoids were cultivated in organoid medium until day 105 counting from the day of transfer into the spinning bottles. Half of the medium was exchanged weekly until week 10 and twice a week afterwards. Two organoids of either control hiPSCs or FOXG1del cells were further analysed after 105 days of cultivation in organoid medium using scRNAseq. In qRT-PCRs single organoids represent the individual n’s and organoids were not pooled.

### Mice

*Foxg1^cre/+^* and *Foxg1^cre/cre^* mice were maintained in a C57Bl/6 background are described by the producers (27), (https://www.jax.org/strain/004337). As multiple recent studies have not shown any discernible sex difference in heterozygote *Foxg1* mouse models (7,13,28,29), in this study, mice were not segregated based on sex. Wildtype mice (C57BL/6J, NMRI or CD1) were obtained from Charles River, Germany or Janvier, France.

### Mouse tissue isolation for RNAseq

Cortex of E13.5 embryos were dissected from Foxg1^cre/cre^ and wildtype mice (Foxg1^+/+^), hippocampi of E18.5 embryos and adult mice were isolated from Foxg1^cre/+^ and wildtype mice (Foxg1^+/+^). Tissue was subsequently used for RNAseq and Immunohistochemistry.

### Mouse hippocampus isolation for primary neuronal culture

Hippocampi of E18.5 embryos and P0 mice were dissected from wildtype mice (C57BL/6J or CD1 with animal license X-17/03S and X-22/02B, Charles River) and collected in 15 ml Hanks’ Balanced Salt Solution (HBSS, Thermo Scientific, USA). The tissue was processed for plating as previously described (20,30). Cells were plated at a density of 100.000 cells/cm^2^ for RNA extraction or 50.000 cells/cm^2^ for microscopy and electrophysiological recordings. Medium change was performed every 72 h by exchanging half of the volume of the medium with fresh NB medium with complements and AraC (1 μg/ml for the first, and 0.5 μg/ml for the second medium change, Sigma/Merck, Germany), cells were harvested at DIV7 (ChIPseq, RIP, RNAseq) (RNAseq) and DIV14 (electrophysiology) based on the performed experiments.

### Production of lentiviral particles and transduction

Lentiviral particles were produced and quantified in HEK293T cells with Mirus TransIT-293 Transfection Reagent (Mirus Bio LLC, Madison, WI, USA) as described before (20). The constructs used are listed in Table 2. Constructs for lentiviral packaging were pMD2.G (Didier Trono, plasmid #12259, Addgene, USA) and psPAX2 (Didier Trono, plasmid #12260, Addgene, USA). Primary hippocampal cells were virally transduced at DIV1 with 1.25 IFU/cell (shRNA-mediated KD as well as rescue with Pantr1 overexpression). Selection of successfully infected cells was performed 72-90 h post-infection (DIV4) by changing half of the volume of the medium with fresh NB medium containing 0.3 μg/ml Puromycin (Sigma/Merck, Germany).

**Table 1:** reagents

| name | catalog number | company | Location |
| --- | --- | --- | --- |
| 2-Mercaptoethanol 50mM | 31350-010 | Gibco, Thermo Scientific | USA |
| Accutase | A1110501 | Gibco, Thermo Scientific | USA |
| Albumin Fraktion V | 8076.4 | Roth | Germany |
| Anti-Adherence Rinsing Solution | 7010 | STEMCELL | Canada |
| AraC | C1768 | Sigma-Aldrich/Merck | Germany |
| B-27 Supplement (50X) 10ml | 17504-044 | Gibco, Thermo Scientific | USA |
| Bradford reagent | 5000006 | Bio-Rad | United Kingdom |
| BSA | 8076.4 | Roth | Germany |
| Chromium Next GEM Single Cell 3' Reagent Kits v3.1 | PN-1000121 | 10X Genomics | CA, USA |
| DAPI | D9542 | Sigma-Aldrich/Merck | Germany |
| DMEM (high glucose) | 11960-044 | Gibco, Thermo Scientific | USA |
| DMEM/NUT.MIX F-12 W/GLUT-I | 31331028 | Thermo Scientific | USA |
| DNAse digestion | #79254 | Qiagen | Germany |
| DPBS | 14190-094 | Gibco | United Kingdom |
| EDTA | AM9260G | Thermo Scientific | USA |
| FBS (Fetal Bovine Serum) | 10500-064 | Gibco, Thermo Scientific | USA |
| Fluorescent Mounting Medium | S3023 | DAKO, Agilent | USA |
| Formaldehyde | 252549 | Sigma-Aldrich/Merck | Germany |
| Glycine | 3908.3 | Roth | Germany |
| GoTaq® qPCR Master Mix | A6002 | Promega | WI, USA |
| Hanks' Balanced Salt Solution (HBSS) | 14025 | Gibco, Thermo Scientific | Germany |
| Hs-PANTR1 | 1199961 | Bio-Techne | Germany |
| Insulin human | 11376497001 | Sigma-Aldrich/Merck | Germany |
| Laminin | L2020-1MG | Sigma-Aldrich/Merck | Germany |
| L-glutamine | 25030081 | Gibco, Thermo Scientific | USA |
| Matrigel | 354277 | Corning | USA |
| MEM NEAA | 11140-035 | Gibco, Thermo Scientific | USA |
| Mirus TransIT-293 Transfection Reagent | MIRUMIR2700 | Mirus Bio LLC | USA |
| Mm-Pantr1 | 483711 | Bio-Techne | Germany |
| mTESR1 medium | #85850 | Stem Cell | Canada |
| N2-Supplement | 17502048 | Gibco, Thermo Scientific | USA |
| NaCl | 92652 | Roth | Germany |
| NEUROBASAL MED SFM | 21103049 | Thermo Fisher | USA |
| Nonidet-40, NP-40 Alternative | 492016 | Calbiochem | USA |
| normal donkey serum | 017-000-121 | Jackson Immuno Research | United Kingdom |
| normal goat serum | G6767 | Sigma-Aldrich/Merck | Germany |
| Not1 | R0189S | New England Biolabs | USA |
| OPTIMEM | 31985-062 | Gibco, Thermo Scientific | USA |
| PageRuler™ Prestained Protein Ladder, 10 to 180 kDa | 26616 | Thermo Scientific | USA |
| Papain Dissociation System | LK003150 | Worthington | USA |
| Penicillin-Streptomycin | 15140122 | Gibco, Thermo Scientific | USA |
| PFA | MC1040051000 | Sigma-Aldrich/Merck | Germany |
| Pierce™ Magnetic RNA-Protein Pull-Down Kit | 20164 | Thermo Scientific | USA |
| Pierce™ RNA 3' End Desthiobiotinylation Kit | 20163 | Thermo Scientific | USA |
| PORN Poly-L- ornithine hydro | P3655-100MG | Sigma-Aldrich/Merck | Germany |
| Primers (see table 3) |  | Sigma-Aldrich/Merck | Germany |
| protease inhibitor | 11873580001 | Sigma-Aldrich/Merck | Germany |
| Protein G Dynabeads | 10004D | Thermo Scientific | USA |
| PSN | 15640-055 | Gibco, Thermo Scientific | USA |
| Puromycin | P8833 | Sigma-Aldrich/Merck | Germany |
| RevertAid Reverse Transcriptase kit | EP0442 | Thermo Scientific | Lithuania |
| RNAeasy Mini Kit | #74104 | Qiagen | Germany |
| RNAscope Fluorescent Multiplex Reagent Kit | 323110 | Bio-Techne | Germany |
| RNAse A | R4642 | Sigma-Aldrich/Merck | Germany |
| SB431542 (50mg) | S1067 | Selleck Chemicals | USA |
| STEMdiff Neural Induction Medium, 250 mL | 5835 | STEMCELL | Canada |
| Stemolecule Dorsomorphin (2mg) | 04-0024 | Reprocell | Japan |
| Sucrose | 4621.1 | Roth | Germany |
| SuperFrost Plus Microscope slides | 11976299 | Thermo Scientific | Germany |
| SuperSignal™ West Femto Maximum Sensitivity Substrate | 34096 | Thermo Scientific | USA |
| T3 MEGAscript kit | AM1338 | Thermo Scientific | USA |
| T4 RNA polymerase | 18005025 | Thermo Scientific | USA |
| Thiazovivin | S1459 | Selleckchem | USA |
| tissue freezing medium | 14020108926 | Leica | USA |
| Trans-blot Turbo Transfer Pack | 1704156 | Bio-Rad | USA |
| TRIS hydrochloride | 90903 | Roth | Germany |
| Triton X | 3051.2 | Roth | Germany |
| Trypsin-EDTA 0,05% | 25300054 | Gibco, Thermo Scientific | USA |
| Tween-20 | P9416 | Merck | Germany |
| with GoTaq® qPCR Master Mix | A6002 | Promega | USA |
| Xho1 | R0146S | New England Biolabs | USA |

**Table 2:**
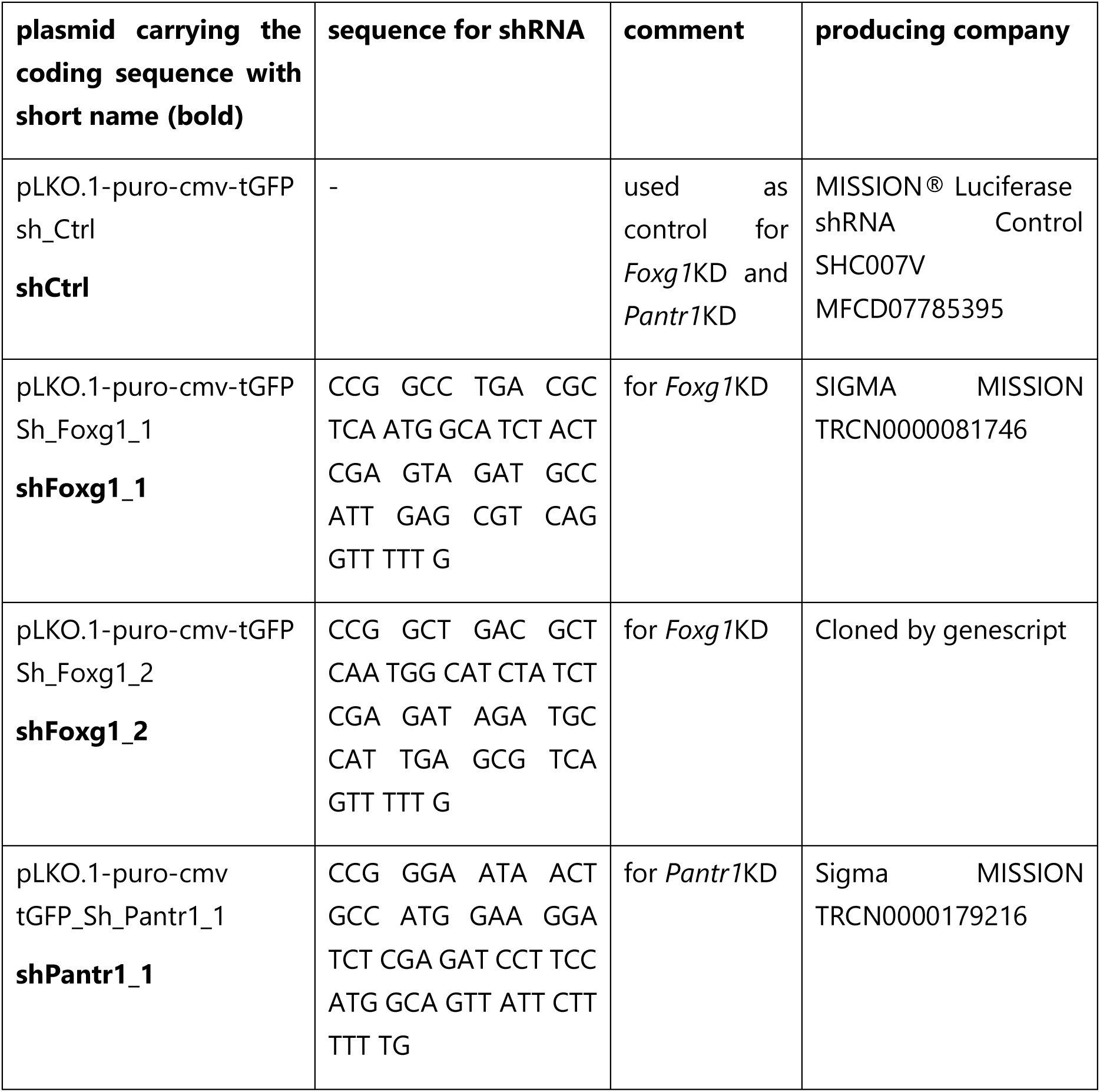
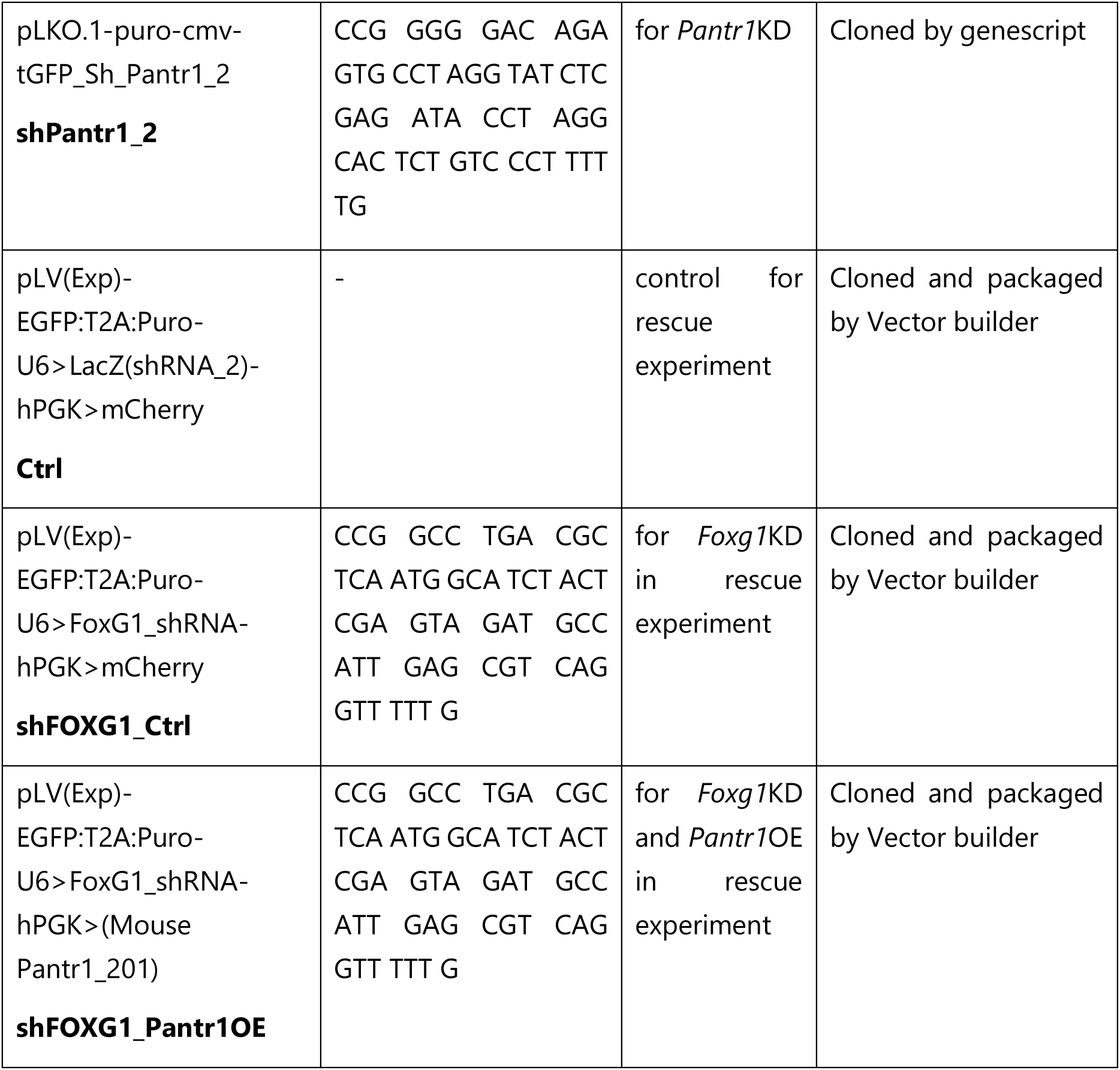
List of plasmids and their shRNA sequence for lentivirus production

### Immunohistochemistry (IHC)

Isolated mouse brains (E18.5 or P0) were fixed for 24 h, human cerebral organoids were fixed for 60 min in 4% PFA (MC1040051000, Merck, Germany). Afterwards the tissues were incubated in 30% sucrose in DPBS (14190-094, Gibco, United Kingdom) overnight at 4°C and were subsequently embedded in tissue freezing medium (Leica, Richmond, USA). Frozen embedded tissues were cut in 14 μm sections and mounted on SuperFrost Plus Microscope slides (Thermo Scientific, Germany). Dried slides were washed two times with DPBS (14190-094, Gibco, United Kingdom), incubated 30 min with 0.25% Triton X (Roth, Germany) in DPBS and 5 min with 0.25% Triton X in DPBS. Blocking was performed in 10% normal donkey serum (017-000-121, Jackson Immuno Research, United Kingdom), 0,1% Triton X in DPBS for 2 h at RT, followed by the incubation with primary antibodies in blocking solution at 4°C overnight. After three washing steps in 0,1% Triton X in DPBS, the cover slips were incubated with secondary antibodies for 1 h at RT (dilution 1:500 in blocking solution), followed by counterstaining with DAPI (1:1000, D9542, Sigma Aldrich, Germany), three additional washing steps with DPBS and mounting in Fluorescent Mounting Medium (DAKO, S3023, Agilent, US). The antibodies used are listed in Table 4.

### Immunocytochemistry (ICC)

For ICC, 50.000 cells plated on coverslips were fixed for 15 min with 4% PFA (MC1040051000, Merck, Germany). Blocking was performed in 10% normal goat serum (G6767, Sigma Aldrich, Germany) and 0,1% Triton X (Roth, Germany) in DPBS (14190-094, Gibco, United Kingdom) for 15 min at RT, followed by the incubation with primary antibodies in blocking solution at 4°C overnight. After three washing steps in DPBS, the cover slips were incubated with secondary antibodies for 1 h at RT, followed by counterstaining with DAPI (1:1000, D9542, Sigma Aldrich, Germany) and mounting in Fluorescent Mounting Medium (DAKO, S3023, Agilent, US). The antibodies used are listed in Table 4.

### Single molecule fluorescent in situ hybridization (smFISH)

For smFISH, cells plated on coverslips were treated following the instructions of manufacturer (RNAscope *Assay for Adherent Cells,* MK50-012 TechNote_09252017, Bio-Techne, Wiesbaden, Germany) for cell fixation. Isolated murine brains (E18.5 or P0) were fixed for 24 h, human cerebral organoids were fixed for 60 min in 4% PFA. Subsequently, the tissues were incubated in 30% sucrose in DPBS (14190-094, Gibco, United Kingdom) overnight at 4°C, followed by embedding in tissue freezing medium (Leica, Richmond, USA). Frozen embedded tissues were cut in 14 μm sections and mounted on SuperFrost Plus Microscope slides (Thermo Scientific, USA). Probe Mm-Pantr1 (murine, 483711, Bio-Techne) or Hs-PANTR1 (human, 1199961, Bio-Techne, Germany) were used following the RNAscope Fluorescent Multiplex Reagent Kit User Manual 320293-USM for detection (Bio-Techne, Wiesbaden, Germany). For smFISH in combination with ICC or IHC, the antibody incubation was performed immediately after smFISH and counterstaining with DAPI was performed at the end before mounting in Fluorescent Mounting Medium (DAKO, S3023, Agilent, US).

### Microscopy and morphometric analysis

DIV7 hippocampal neurons stained with MAP2 (1:200, ab32454, abcam, United Kingdom) were imaged with the microscope Axio Imager M2, objective EC Plan NeoFluar 40x/0.75 M27, Axiocam 506, software ZEN 3.0. Dendrites were traced manually using the Sholl-analysis plugin for ImageJ 1.52p by a second experimenter blindly. Dendrite complexity was assessed by measuring the dendritic intersections in concentric circles per 10 μm from the cell soma and significance was determined using two-way ANOVA test.

For dendritic spine density analysis, biocytin-filled neurons were stained with Alexa Fluor™ 633-conjugat (1:500, S21375, Life Technologies, USA), Z-stack pictures were acquired using SP8 confocal microscope, objective HC PL APO CS2 63x/1.40 OIL, zoom 2x and resolution 2048 x 2048. The analysis was performed by counting spines on 20 μm length dendrite, 3 dendrites for each neuron were quantified using LAS X 3.5.2.18963.

For localization and co-localization studies Z-stack pictures were acquired using SP8 confocal microscope, objective Zeiss HC PL APO CS2 63x/1.40 OIL, pixel size 0.361 x 0.361 µm and analysis was performed with LAS X 3.5.2.18963 software (Leica Microsystems, Wetzlar, Germany). For co-localization of *Pantr1* and FOXG1 additional to the Leica system, images were taken with the 60x objective of the BC43 (Oxford Instruments Andor, United Kingdom) and processed with Imaris 10 (Oxford Instruments, United Kingdom).

### Co-immunoprecipitation of protein and RNA

Co-Immunoprecipitation (Co-IP) was performed as previously described (22). In brief, P0 whole brain tissue or DIV7 cells weres lysed in lysis buffer (300 mM NaCl, 20 mM Tris, 1 mM EDTA, 0.5% Nonidet-40 (NP-40), pH 7.4) supplemented with protease inhibitor (Sigma-Aldrich/Merck, Germany) and incubated for 30 min on ice, triturating every 10 min 20 times. After a 10 min centrifugation at 13000 rpm, the supernatant was collected. The samples were diluted dropwise to reach a salt concentration of 100 mM and NP-40 concentration of 0.15% using equilibration buffer (20 mM Tris, 1 mM EDTA, pH 7.4). Protein concentrations were determined with Bradford reagent (Bio-Rad, United Kingdom). 10% input was saved and equal amounts of protein were used for all Co-IPs. Protein G Dynabeads (10004D, Thermo Scientific, USA) were washed with Co-IP buffer (100 mM NaCl, 20 mM Tris, 1 mM EDTA, 0.15% NP-40 pH 7.4), and incubated with Co-IP antibodies at 4°C overnight. The following antibodies were used: anti-FOXG1 (#61211, Active Motif, USA), IgG (#C15410206 or #C15400001, Diagenode, Belgium). Antibody-coupled beads were washed once with Co-IP buffer before Co-IP. Cell lysates were precleared with Protein G Dynabeads for 1 h at 4°C, subsequently transferred to antibody-coupled beads, and incubated overnight at 4°C in rotation. If the aim was to investigate binding and complex forming in absence of RNA, RNAse A (1:400, R4642, Sigma-Aldrich/Merck) was added to this overnight step. Antibody-coupled beads were washed 3 times with Co-IP buffer. Depending on downstream analysis they were either resuspended in 1X Laemmli sample buffer (protein detection by immunoblotting) or in RLT buffer of the RNA isolation kit (RNA detection by native RIPseq or RIP-qRT-PCR). Protein-antibody complexes were eluted by incubating the beads at 70°C for 10 min at 550 rpm. 10% input and the complete Co-IP sample were used for immunoblotting.

### Immunoblotting (IB)

Co-immunoprecipitation (Co-IP) samples were loaded onto 10% SDS-polyacrylamide gels and electrophoresed at 120 V for 1.5 h in Tris-Glycine running buffer. Subsequently, proteins were transferred onto PVDF membranes (Trans-blot Turbo Transfer Pack, Bio-Rad, USA) using the Trans-blot Turbo Transfer System (Bio-Rad, USA), following the manufacturer’s instructions. Afterwards, the membranes were blocked with 5% BSA (Roth, Germany) in Tris-buffered saline (TBS) for 1 h. For immunoblotting, the following antibodies were utilized: anti-FOXG1 (anti-FOXG1, #61211, Active Motif, Carlsbad, CA, USA), diluted in 5% BSA in TBST (Tris-buffered saline with 0.1% Tween-20 (Merck, Germany)). After three washes with TBST, the membranes were incubated with Tidyblot (diluted 1:5000 in 5% BSA in TBST, Biorad, USA), for 1 h. Following two washes with TBST and two with TBS, the membranes were developed using SuperSignal™ West Femto Maximum Sensitivity Substrate (Thermo Scientific, USA) and imaged with the LAS ImageQuant System (GE Healthcare, USA).

### RNA isolation, reverse transcription, and quantitative real-time PCR analysis

RNA was isolated from tissue, organoids, cell samples or coIP beads with the RNAeasy Mini Kit (#74104, Qiagen, Germany) including an on-column DNAse digestion (#79254, Qiagen, Germany). The isolated RNA was either used for further RNAseq or for quantitative real-time PCR (qRT-PCR) analysis. For qRT-PCR analysis, we harvested 1 Mio cells or one organoid in RLT buffer (#74104, Qiagen, Germany). cDNA was synthesized with the RevertAid Reverse Transcriptase kit (EP0442, Thermo Scientific, Lithuania) and subsequently used for a qRT-PCR with GoTaq® qPCR Master Mix (A6002, Promega, WI, USA) following the manufacturer’s protocol. Primers (Sigma/Merck, Germany) are listed in Table 3. qRT-PCR after coIP was performed with three technical replicates and the ct values were used to calculate the percentage of RNA left after co-IP compared to the cell lysate (input). All other analyses were performed as described before (20). Data are presented as mean ± SEM. P-values were calculated using unpaired, two tailed Student’s t-test or one-way ANOVA: *p < 0.05, **p < 0.01, ***p < 0.001 with GraphPad Prism 6.

**Table 3:**
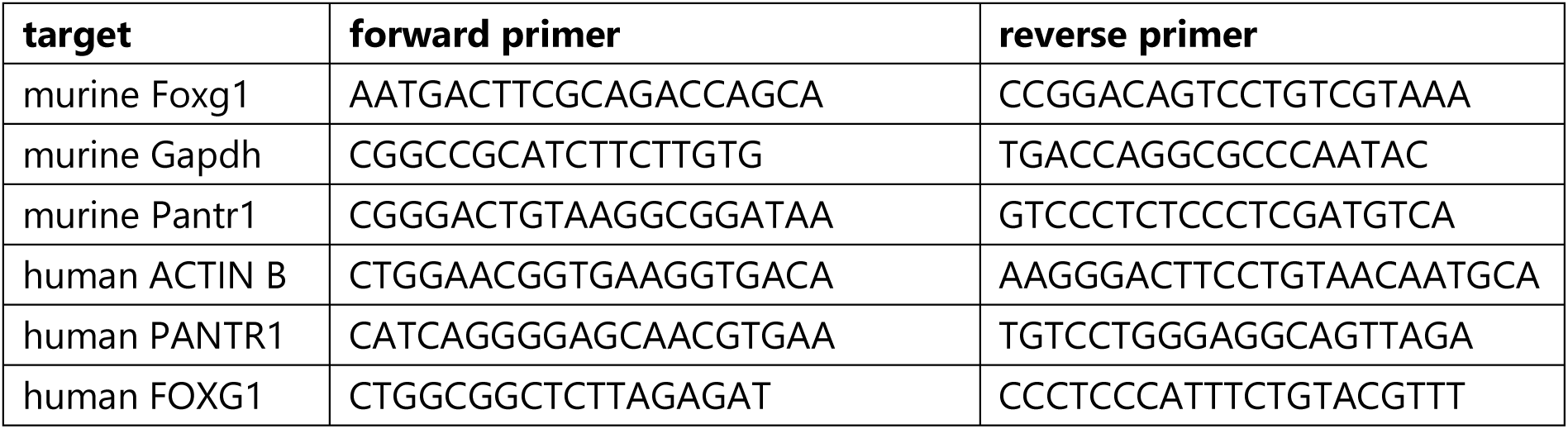
List of primers for qRT-PCR

**Table 4:** List of antibodies

| name | usage and dilution | source |
| --- | --- | --- |
| FOXG1 | IHC, 1:500 | Abcam ab196868, United Kingdom |
| FOXG1 | IB, 1:1000, coIP, RIP | active motif #61211, USA |
| GFAP | IHC, 1:500 | life technologies 13-0300, USA |
| MAP2 | ICC, 1:200 | Abcam ab32454, United Kingdom |
| NeuN | IHC, 1:150 | Abcam ab177487, United Kingdom |
| PAX6 | IHC, 1:150 | BioLegend 901301, USA |
| TBR1 | IHC, 1:200 | Abcam ab31940, United Kingdom |
| TBR2 | IHC, 1:200 | Invitrogen 14-4875-82, USA |
| TUJ-1 (TUBB3) | IHC, 1:200 | BioLegend MMS-435P-100, USA |
| vGLUT1 | IHC, 1:500 | Synaptic Systems 135 303, Germany |
| vGLUT2 | IHC, 1:500 | Abcam ab79157, United Kingdom |
| anti Mouse-Alexa488 | ICC, IHC 1:500 | Dianova (715-545-151), Germany |
| anti Mouse-Alexa594 | ICC, IHC 1:500 | Dianova (715-585-151), Germany |
| anti Rabbit-Alexa488 | ICC, IHC 1:500 | Dianova (711-545-152), Germany |
| anti Rabbit-Alexa594 | ICC, IHC 1:500 | Dianova (711-585-152), Germany |
| anti Rat-Alexa488 | ICH, 1:500 | ThermoFisher A21208, USA |
| Streptavidin-Alexa-633 | ICC, 1:500 | Life Technologies S21375, USA |
| Tidyblot | IB 1:5000 | Bio-Rad Tidyblot STAR209P, USA |

### Single cell (sc)RNAseq Library Preparation

Two organoids of each line were collected at day 105 and single cells were obtained according to the protocol of the Papain Dissociation System (Worthington, Lakewood, USA), with centrifugation steps carried out at 4°C. Library preparation was performed with the 10x Genomics kit according to the manufacturer’s instructions (Chromium Next GEM Single Cell 3’ Reagent Kits v3.1, 10X Genomics, CA, USA).

### Bulk RNA-sequencing and RIPseq

Total RNA extracted from mouse hippocampi, cortex or DIV7 primary hippocampal neurons were used for RNAseq. Total RNA extract of FOXG1- and IgG-coIP respectively were used for RIPseq. The cDNA libraries were constructed using TruSeq total RNA sample preparation kit (Illumina, USA), following the manufacturer’s instructions and sequenced on Illumina HiSeq 3000 as paired-end 101 bp reads. For RNAseq, the procedure included depletion of rRNA prior to double-stranded cDNA synthesis and library preparation. Each replicate (n) came from independent cell cultures of different litters respectively from independent mice.

### ChIPsequencing

For FOXG1-ChIPseq analysis, wildtype or virally transduced primary hippocampal neurons were fixed and collected as previously described (22), following the instructions of the servicing company (Active Motif, Carlsbad, CA, USA). For FOXG1-ChIPseq analyses after *Pantr1*KD, primary hippocampal neurons transduced with lentiviral shPantr1_1 constructs or control constructs were fixed on DIV7. The FOXG1-ChIPseq of cortical organoids was performed after their fixation using the same protocol as for cells.

A custom Illumina library type on an automated system (Apollo 342, Wafergen Biosystems/Takara) was used to generate ChIPseq libraries, which were subsequently sequenced on Illumina NovaSeq 6000 as single-end 75 bp reads (Active Motif, Carlsbad, CA, USA). Antibodies used for the immunoprecipitation were either anti-FOXG1 (#61211, Active Motif, Carlsbad, CA, USA) for murine samples or anti-FOXG1 (ab18259, abcam, United Kingdom) for human samples.

### RNA pulldown of adult hippocampus protein extracts

*Pantr1* lincRNA was cloned into pBluescript plasmid such that the sense strand was in frame with T7 promoter and antisense strand with T3. *Pantr1* RNA sense strand was synthesized *in vitro* using linear pBluescript-Pantr1 plasmid (linearized using Xho1 (New England Biolabs, USA)) and T7 RNA polymerase (T7 MEGAscript kit, Thermo Scientific, USA) as per the manufacturer’s protocol. Similarly, *Pantr1* antisense strand was synthesized by using Not1 (New England Biolabs, USA) digested linear pBluescript-Pantr1 plasmid and T4 RNA polymerase (T3 MEGAscript kit, Thermo Scientific, USA). Next, 10 µg of *Pantr1* sense RNA were used to label the 3’ end using Pierce™ RNA 3’ End Desthiobiotinylation Kit (Thermo Scientific). After labelling, *Pantr1* sense RNA was purified to remove the unincorporated labelled nucleotides and RNA pull down was performed using Pierce™ Magnetic RNA-Protein Pull-Down Kit (Thermo Scientific, USA), according to the instructions. Protein extracted from six week old NMRI mouse hippocampus were used for the RNA pull down assay (with animal license X-14/04H). RNA pulldown samples (n=3) were collected and subjected to LC-MS/MS to identify the proteins interactome of *Pantr1* lincRNA.

### Bioinformatics

#### Single cell RNAsequencing analysis

Reads from single cell (sc)RNAseq were aligned to the GRCh38 human reference genome, and the cell-by-gene count matrices were produced using the RNA STARSolo on the Galaxy Platform. Data were analysed using the Seurat R package v.4.3.0 with R v.4.2.2. The following parameters were used for filtering cells: cut_nUMI <-500; cut_nGene <-250; max_nGene <-8500; cut_log10GenesPerUMI <-0.80; cut_mito.Ratio <-0.2; cut_ribo.Ratio <-0.2.

The expression values were then normalised using the ’SCTransform v2’ function. Subsequently, Principal Component Analysis (PCA) was applied to highly variable genes identified by the ’FindVariableGenes’ function, retaining the top 40 principal components (PC) based on Seurat’s Elbow Plots and determination of percent variation associated with each PC. Clustering was then executed on the PCA-reduced data utilising the ’FindNeighbors’ and ’FindClusters’ functions, with a resolution set at 0.4. Uniform Manifold Approximation and Projection (UMAP) was employed for dimensionality reduction, providing a visual representation of the data. For cell type annotation, differential expression analysis was conducted between clusters via the ’FindAllMarkers’ function, considering genes with a log-fold change greater than 0.1 and an adjusted p-value less than 0.05 as differentially expressed. FOXG1 and *PANTR1* expression levels were plotted using VlnPlot and FeaturePlot functions within the Seurat package. Cell type proportions were calculated using scProportionTest tool with default settings (31).

#### Bulk RNA-, ChIP-, RIPsequencing analysis

The Galaxy platform as well as R based analysis were used to analyse the RNAseq, FOXG1 ChIPseq and RIPseq data. Mapping was performed on mouse genome build mm10 (GRCm38) or human genome build hg38 depending on organism.

Quality control, trimming, mapping of the RNA sequencing fastq files, and generation of gene-level counts was done on the Galaxy platform (32). Differential expression analysis of DIV7 hippocampal neurons (Fig. 5B) was done using DESeq2 (1.34.0) on R (4.2.0) on the count matrix output from featurecounts (33). GO term enrichment analyses were done using clusterProfiler (4.2.2) (34). Visualisations of volcano plots and heatmaps were done using EnhancedVolcano (1.12.0) and pheatmap (1.0.12) packages, respectively (35-37).

Differential expression analysis for RNAseq of the adult and E18.5 hippocampus sample (Fig. 3C) was done using DESeq2 (v. 1.22.1) (33) on count matrices output from snakePipes (featureCounts, v. 1.6.4) (38). A linear model controlling for batch effects (e.g., ∼batch + treatment or ∼ batch + condition) was used and apeglm log2(Fold Change) shrinkage was applied.

RIPseq fastq files were processed on the Galaxy platform. Trimmed and quality filtered reads were aligned to mm10 genome assembly using RNAStar (2.7.10b). FOXG1-coIP reads were normalized to IgG-coIP reads using bamcompare (v. 3.5.4) and the average calculated using bigwigAverage (v. 3.5.4). Differentially immunoprecipitated RNA was calculated using DESeq2 (v. 2.11.4.0.8) after generating count matrices (featureCounts, v. 2.0.3).

FOXG1 ChIPseq quality controls (FastQC) were performed on the galaxy server (usegalaxy.eu). The sequences were mapped against mm10 with Bowtie and Peaks were called using MACS callpeak. The coverage was calculated followed the analyses as described previously (22).

GO enrichment and differential GO-term analyses were performed using clusterProfiler (v. 4.2.2) (34).

Motif enrichment and differential motif enrichment analyses of the ChIPseq dataset were done using gimmemotifs on Python (39).

Visualisations were done in Galaxy, R (v. 4.1) and Python (v. 3.6). Heatmaps were plotted using heatmap2 (Galaxy Version 3.0.1), genome tracks using pyGenomeTracks (Galaxy Version 3.6), venn diagrams using ggVenn (Linlin Yan (2021), ggvenn: Draw Venn Diagram by “ggplot2”. R package version 0.1.9) and VennDiagram (Hanbo Chen, VennDiagram: Generate High-Resolution Venn and Euler Plots. R package version 1.7.3. (2022) packages, volcano plots using EnhancedVolcano (BioConductor, v. 3.13) (B. K. Lewis M Rana S., EnhancedVolcano: Publication-ready volcano plots with enhanced colouring and labeling. R package version 1.14.0, (2022)), violin plots using ggplot2 (v. 3.3.5) (40).

### Statistics

GraphPad Prism software (version 6) was used for statistical analyses. Depending on the experimental design, two-sided unpaired Student’s t-test or two sided one-way ANOVA followed by Tukey multiple comparisons were used for the analyses of qRT-PCR experiments. RNAseq and RIPseq counted features were statistically analysed using the Deseq2 tool (33) provided by the Galaxy platform (37). Statistics in scRNAseq and pseudobulk RNAseq analysis of the organoids are described in the bioinformatics section in the supplement. Statistical significance in ChIPseq data were analysed using the Diffbind tool (41) on the Galaxy platform. Electrophysiological recordings were tested using Mann-Whitney test, and Sholl analysis was tested with two-sided two-way ANOVA followed by Sidak’s multiple comparison test.

### Reagents

## RESULTS

### iPSC-derived human brain organoids as model for FOXG1-syndrome

To explore the molecular underpinnings of FOXG1-syndrome in human model systems, we used human patient derived iPSC carrying one allele with a full deletion of FOXG1 (FOXG1del). We generated hCO, which we cultured for 105 days (d). hCO expressed marker genes for progenitors, glutamatergic and GABAergic neuron populations (**Fig. S1A, B**), including FOXG1, in the healthy donor. FOXG1del derived hCO expressed less FOXG1, as expected (**Fig. S1B**). To characterise the hCO at the transcriptional level, we performed single cell (sc) RNAseq on 11258 cells from the healthy donor (HD) control and 9292 cells from the FOXG1del organoids. UMAP representation of cell clusters retrieved stem and progenitor cells, as well as diverse stages and subtypes of excitatory and inhibitory neurons (**Fig. 1A**). By plotting *FOXG1* expression levels, we observed broad expression of *FOXG1*, and reduced levels of it upon *FOXG1* deletion compared to the HD (**Fig. 1B**). Cluster-wise comparison of expression levels of *FOXG1* in HD and FOXG1del hCO revealed that *FOXG1* was transcribed by all cells but in different levels (**Fig. 1C**). Upon deletion of one *FOXG1* allele, all cell clusters showed reduced *FOXG1* transcription (**Fig. 1C**). We observed significant changes in cell proportions for some specific clusters upon *FOXG1* deletion, including apical progenitor cells (APC), truncated radial glia (tRG), three interneuron populations (SST+IN, nVP, IN-LGE) and two excitatory neuron populations (DL-EN, early UL-EN) (**Fig. 1D**). qRT-PCR using the entire hCO (bulk) for RNA extraction confirmed significantly decreased *FOXG1* expression in FOXG1del hCO (**Fig. 1E**). Functional GO-terms of pseudobulk RNAseq analysis associated lower expressed genes in FOXG1del hCO to processes like regulation of neurogenesis and neuron fate commitment, whereas higher expressed genes correlated to processes like nerve development and neuron apoptosis. Genes related to telencephalon and forebrain development as well as axonogenesis and synapse organisation presented in both clusters of lower and higher expressed genes (**Fig. 1F**).

**Fig. 1:**
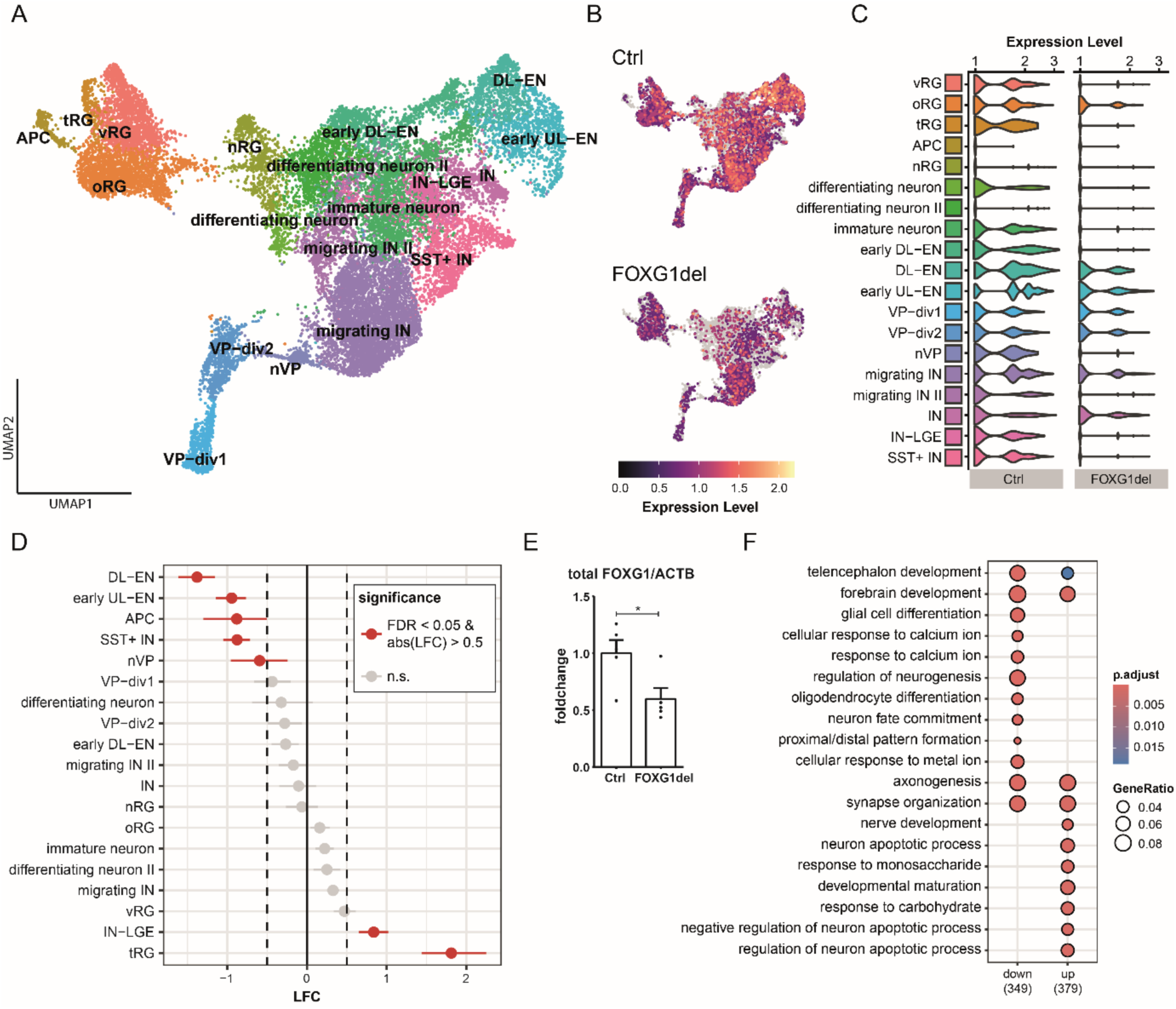
iPSC-derived hCO model FOXG1-syndrome. **(A)** UMAP plots, colour-coded and annotated by cell type, depicting the d105 cerebral organoids (hCO, brain organoids) obtained from a healthy donor and a FOXG1-syndrome patient featuring a heterozygous deletion of *FOXG1* (FOXG1del). n=2. Number of cells: Control=11258, FOXG1del=9292. Abbreviations: APC: Astrocyte progenitor cells, DL-EN: Deep layer excitatory neuron, IN: Interneuron, IN-LGE: interneurons of the lateral ganglionic eminence, nIP: neuronal Intermediate Progenitor, nRG: neuronal radial glia (RG), nVP: neurogenic ventral progenitor, oRG: outer RG, SST+IN: Somatostatin-positive Interneurons, tRG: truncated RG, UL-EN: Upper layer excitatory neuron, VP-div: Ventral Progenitor-dividing, vRG: ventricular RG. **(B)** UMAPs of healthy and FOXG1del organoids coloured by expression level of *FOXG1* (low: grey; high: purple to yellow). **(C)** Violin plots of cell-type specific expression distribution of *FOXG1*. *FOXG1* expression reduced upon *FOXG1* deletion in all cell populations. **(D)** Dotplot of the ratio of cell type proportions of FOXG1del organoids compared to the control organoids. Shown is the log_2_(foldchange) (LFC). A significance threshold of false discovery rate (FDR) < 0.05 and a LFC < -0.5 or > 0.5 was applied. Significant values are marked in red. FOXG1del hCO present lower numbers of SST+ IN, APC, early UL−EN, DL−EN and nVP, whereas numbers of IN−LGE and tRG are significantly increased. **(E)** RT-qPCR showing expression levels of *FOXG1* (with the housekeeping gene ACTIN B (*ACTB*) as reference). The fold change in expression in FOXG1 deletion (FOXG1del) hCO is compared to that in control organoids. n=5. **(F)** GO-term enrichment analysis of pseudobulk DEGs upon deletion of one *FOXG1* allele split according to DEGs with either increased or decreased expression compared to healthy donor hCOs. Given are the top enriched GO terms with a FDR > 0.5.

As FOXG1 functions at the chromatin level, we performed ChIPseq using FOXG1 antibody at d105 of the HD hCO. FOXG1 bound TSS distributed similar to mouse (42), i.e. concentrated around the TSS (cluster 3, **Fig. 2A**) but also dispersed (cluster 1, **Fig. 2A**). Similar to what has been described for the mouse hippocampus, FOXG1 bound mainly intronic and intergenic regions as well as promoters (**Fig. 2B**). Genes bound by FOXG1 in HD conditions classified as genes involved in axonogenesis and forebrain and telencephalon development (cluster 1) as well as ncRNA processing and RNA splicing (cluster 3) in enriched functional GO-terms (**Fig. 2C**). Intersection of differentially expressed and FOXG1 bound regions revealed 266 potentially direct FOXG1 target genes in hCO that changed expression upon deletion of one *FOXG1* allele (**Fig. 2D**). Functional GO-terms associated the differently expressed genes in FOXG1del hCO to processes like forebrain development, axonogenesis and axon guidance (**Fig. 2E**). Together, the characterisation of HD and FOXG1del hCO, indicated similarities to the mouse model. We concluded that parallel analysis of human and mouse multiomic data might help identifying and functionally assessing regulative factors of FOXG1. Here, we further exploited whether lncRNA are part of the FOXG1-regulative network.

**Fig. 2:**
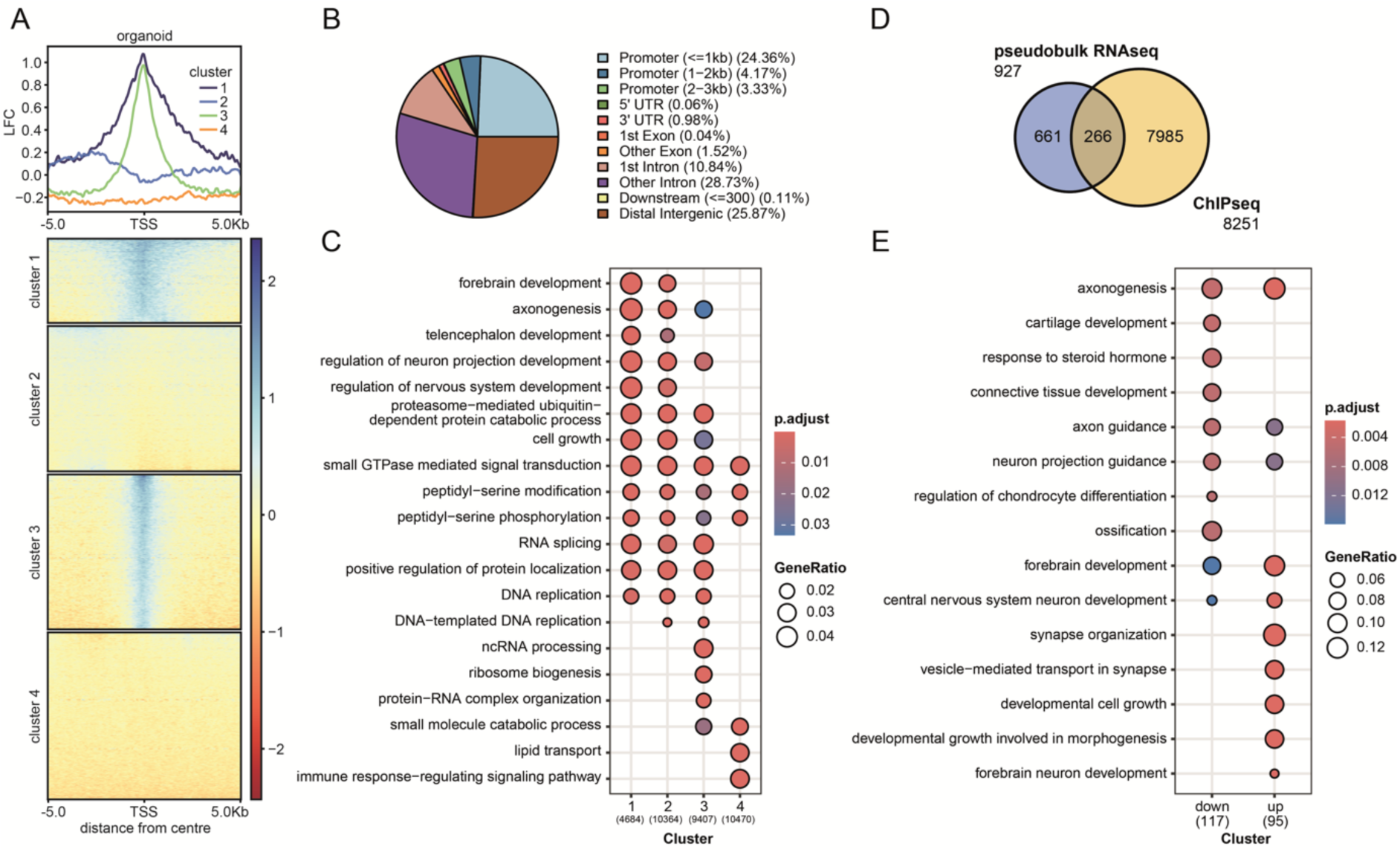
FOXG1 binds to gene loci encoding for genes in hCO correlated with neural development and RNA processing. **(A)** Heatmap of differently bound regions (DBRs) of FOXG1, 5 Kb up-/downstream of FOXG1 peak summits in d105 heathy donor hCO. The metaprofiles (top) show the mean log2(foldchange) (LFC) of 4 k means clustered regions compared to the input. **(B)** Genome-wide distribution of bound regions of FOXG1 in d105 healthy donor hCO in percentage for near and far promotor regions, 5’UTR, intronic and exonic as well as distal intergenic regions. **(C)** GO-term enrichment analysis of genes located near to the peaks in the ChIPseq data split according to the clustering in Fig 2A. Given are the top enriched GO terms with a FDR > 0.5. **(D)** Venn diagram of FOXG1 enriched peaks and pseudobulk DEGs upon deletion of one FOXG1 allele (FOXG1del) in d105 hCO. **(E)** GO-term enrichment analysis of common genes in the ChIPseq and pseudobulk RNAseq data sets split according to DEGs with either increased or decreased expression compared to healthy donor d105 hCO. Given are the top enriched GO terms with a FDR > 0.5.

### FOXG1 binds to gene loci encoding for lncRNAs in mouse and human

We explored both mouse and human ChIPseq as well as transcriptomic data sets for the presence of lncRNAs to identify their potential role in the context of FOXG1-syndrome. For exploring whether FOXG1 would directly regulate l(i)ncRNA expression in the mouse, we identified all loci in close vicinity or within l(i)ncRNA-encoding genes from ChIPseq experiments of adult hippocampus (*in vivo*) as well as primary hippocampal neurons (*in vitro*) (**Fig. 3A**). We identified 520 genomic loci associated with l(i)ncRNAs *in vivo*, and 788 *in vitro*, of which 488 were shared among both data sets (**Fig. 3B**). To investigate whether l(i)ncRNAs are affected upon loss of FOXG1, we utilised *Foxg1^cre/+^*adult hippocampus, the developing *Foxg1^cre/+^* E18.5 hippocampus (22), and the *Foxg1^cre/cre^* E13.5 cerebral cortex RNAseq datasets. Our RNAseq analyses revealed altered expression of 8 l(i)ncRNAs in adult hippocampal tissue, 17 in the developing hippocampus, and 193 in the cerebral cortex upon *Foxg1* mutation, using a threshold of adjusted p-value (padj) < 0.05 (**Fig. 3C, D**). L(i)ncRNAs common to all mouse transcriptome data sets were *Pantr1* and *Firre*, of which only *Pantr1* occurred in the shared targets of *in vivo* and *in vitro* FOXG1 ChIP peaks (**Fig. 3B**). In support of these findings, we validated decreased levels of *Pantr1* in the *Foxg1^cre/+^* adult hippocampus using RT-qPCR (**Fig. 3E**).

**Fig. 3:**
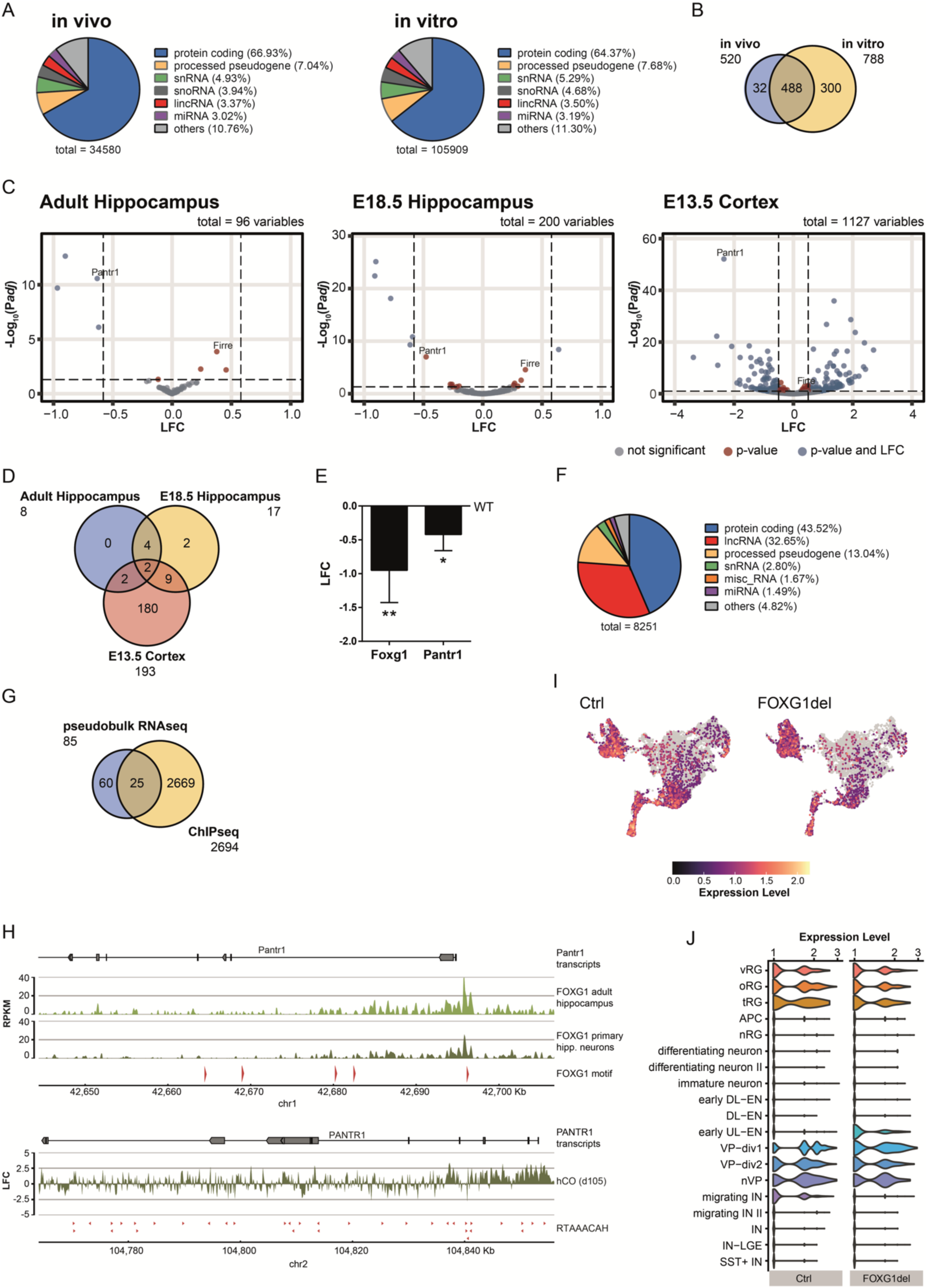
*Pantr1* is a direct FOXG1 target in mouse and human. **(A)** Pie chart of the distribution of FOXG1 peaks regarding different subtypes of transcripts, derived from *in vivo* (adult, hippocampus) or *in vitro* (E18.5, cultivated hippocampal neurons) mouse ChIPseq data (GSE189119 (22)). **(B)** Venn diagram of FOXG1 bound regions annotated as l(i)ncRNAs *in vivo* and *in vitro*, indicating an overlap of 488 between both conditions. **(C)** Volcano plots of differentially expressed l(i)ncRNAs upon deletion of *Foxg1* in adult hippocampal tissue (*Foxg1^cre/+^* vs wildtype mice, n=2; GSE189119), E18.5 hippocampal tissue (*Foxg1^cre/+^* vs wildtype mice, n=3, GSE189119), and E13.5 cortical tissue (*Foxg1^cre/cre^* vs wildtype mice, n=5; GSE95833). The log_2_(foldchange) (LFC) compared to the control is shown on the x-axis while the y-axis shows the mean expression value of -log10 (p-value). Indicated are the threshold at adjusted p-value (padj) of 0.05 and an LFC of +/-0.58 with dashed lines. DEG fulfilling both conditions are coloured in blue, genes with a padj higher than 0.05 but an LFC between -0.58 and 0.58 are red and all other genes grey. **(D)** Venn diagram of differentially expressed l(i)ncRNAs upon *Foxg1* deletion in adult hippocampal tissue (*Foxg1^cre/+^* vs wildtype mice), E18.5 hippocampal tissue (*Foxg1^cre/+^* vs wildtype mice), and E13.5 cortical tissue (*Foxg1^cre/cre^* vs wildtype mice). **(E)** *Foxg1* and *Pantr1* expression in *Foxg1^cre/+^* adult hippocampus tissue compared to wildtype mice (WT). Shown is the log_2_(foldchange) (LFC). *Gapdh* was used as housekeeping gene. Results of qRT-PCRs are shown as mean ± SEM. P-values were calculated using unpaired, two tailed Student’s t-test: *p < 0.05, **p < 0.01. n=3. **(F)** Pie chart of the distribution of FOXG1 peaks regarding different subtypes of transcripts, derived from hCO ChIPseq data (Fig. 2A and B). **(G)** Venn diagram of differentially expressed l(i)ncRNAs in FOXG1del compared to HD hCO (pseudobulk RNAseq) and FOXG1 bound regions annotated as l(i)ncRNA as revealed by ChIPseq data from d105 hCO (Fig. 2A and B). **(H)** Top: ChIPseq of FOXG1 in wildtype *in vivo* adult hippocampus and *in vitro* E18.5 mouse primary hippocampal neurons (DIV11) (22) shows FOXG1 bound regions at the *Pantr1* gene loci. The upper track shows an overlay of all *Pantr1* transcript variants in this region. On the x-axis the position on chromosome 1 (chr1) is indicated in Kb of the mm10 genome. Mapped tracks show the reads per kilobase of transcript per million mapped reads (RPKM) for each condition. Lower part shows all potential FOXG1 binding sites containing the mouse FOXG1 motifs. Bottom: ChIPseq of FOXG1 from d105 hCO shows FOXG1 bound regions at the *Pantr1* gene loci. The log_2_(foldchange) (LFC) compared to the input is shown. On the x-axis the position on chromosome 2 (chr2) is indicated in Kb of the hg38 genome. Lower part shows all potential FOXG1 binding sites containing the motif RTAAACAH. **(I)** UMAPs of healthy and FOXG1del organoids coloured by expression level of *PANTR1* (low: grey; high: purple to yellow). **(J)** Violin plots of cell-type specific expression distribution of *PANTR1*. *PANTR1* expression reduced significantly upon *FOXG1* deletion in different radial glia highlighted in brownish colour, and in the GABAergic lineage of IN progenitors, migrating IN, and early IN, as highlighted in blue.

In hCO, we identified 2694 genomic loci related to l(i)ncRNAs bound by FOXG1 (**Fig. 3F**). Pseudobulk RNAseq analysis of the hCO transcriptome revealed 85 l(i)ncRNAs among the DEGs, comparing FOXG1del to HD (**Fig. 3G**). 25 l(i)ncRNA were common in both ChIP and pseudobulk RNAseq datasets. The only common l(i)ncRNA between mouse and human datasets, analysed in the same manner, was *PANTR1*. In ChIPseq tracks for both human and mouse, *PANTR1* loci showed multiple potential FOXG1 binding sites, a subset of which showed enrichment of FOXG1 (**Fig. 3H**). The potential binding site in hCO was assigned to cluster 1 (**Fig. 2A**). This data together rendered *PANTR1* as direct target gene of FOXG1.

Next, we plotted *PANTR1* expression levels in hCO in single cell resolution, and observed reduced levels of *PANTR1* upon *FOXG1* deletion compared to the HD (**Fig. 3I**). Cluster-wise comparison of expression levels of *PANTR1* in HD and FOXG1del hCO revealed that *PANTR1* expression was more cell-type restricted compared to *FOXG1* expression (**Fig. 3J, 1C**). *PANTR1* was mainly expressed by distinct radial glia cells and developing interneurons. Accordingly, upon *FOXG1* deletion, *PANTR1* expression reduced in radial glia and interneurons. Interestingly, two cell clusters seemingly showed increased expression of *PANTR1*, namely early upper layer neurons and neurogenic ventral progenitor cluster (nVP) (**Fig. 3J**). We observed changes in cell proportions for some clusters upon *FOXG1* deletion (**Fig. 1D**). However, these changes did not correlate clearly with the expression changes for *PANTR1*. Summarising, *PANTR1* expression de- and increased depending on the cell population in FOXG1del compared to HD.

### FOXG1 associates with the lncRNA *Pantr1*

Members of the forkhead (fkh) domain family bind to RNA (43), and one recent report shows that L1 mRNA is not only a transcriptional target of FOXG1, but also associated to FOXG1 (44). Given these indications for FOXG1 complexing with RNA, we applied RNA immunoprecipitation (RIP) for FOXG1 followed by qRT-PCR for *Pantr1* using mouse wildtype P0 brains. This experiment identified *Pantr1* as a transcript among the precipitated RNAs (**Fig. 4A**). Additionally, we performed RIPseq for FOXG1 and detected several RNAs that enriched in the FOXG1 RIP compared to the IgG control (**Fig. 4B**). Upon RIPseq, however, *Pantr1* was not among the top enriched RNAs, but was present with only few reads compared to the most enriched RNA, *Map1b* (**Fig. S2A, B**). For the reverse viewpoint, we used mass spectrometry of hippocampal proteins precipitated through biotinylated *Pantr1*, which enriched a diverse set of proteins. However, we did not detect FOXG1 among them (**Fig. 4C**). Analysis of FOXG1 interacting proteins (23) revealed overlapping FOXG1 interactors and *Pantr1* binding proteins (**Fig. 4D**). These findings strongly suggested an association between FOXG1 and *Pantr1*, although it may occur directly or be bridged by another protein. To further characterise a functional association between FOXG1 and *Pantr1*, we immunoprecipitated FOXG1 protein upon *Pantr1*KD. We detected FOXG1 in several bands, with molecular weights around 50, 65 and 70 kDa, of which the 65 kDa signal was undetectable upon *Pantr1*KD (**Fig. 4E**). This finding suggested a *Pantr1*-mediated occurrence of a specific FOXG1^65^ isoform that coexisted with the monomeric FOXG1^50^. The FOXG1^65^ isoform could contain RNA, represent a specific, RNA-dependent conformation, or might be a heterodimer with a small 10-15 kDa protein. For assessing potential colocalisation *in vivo*, we combined *PANTR1* smFISH and FOXG1 immunostaining in hCO and mouse cortical sections, and observed that the signals were close to each other within the nucleus in human and mouse cells (**Fig. 4F, G, S3A, B**). Together, the findings indicate that FOXG1 and *PANTR1* likely acted within one pathway.

**Fig. 4:**
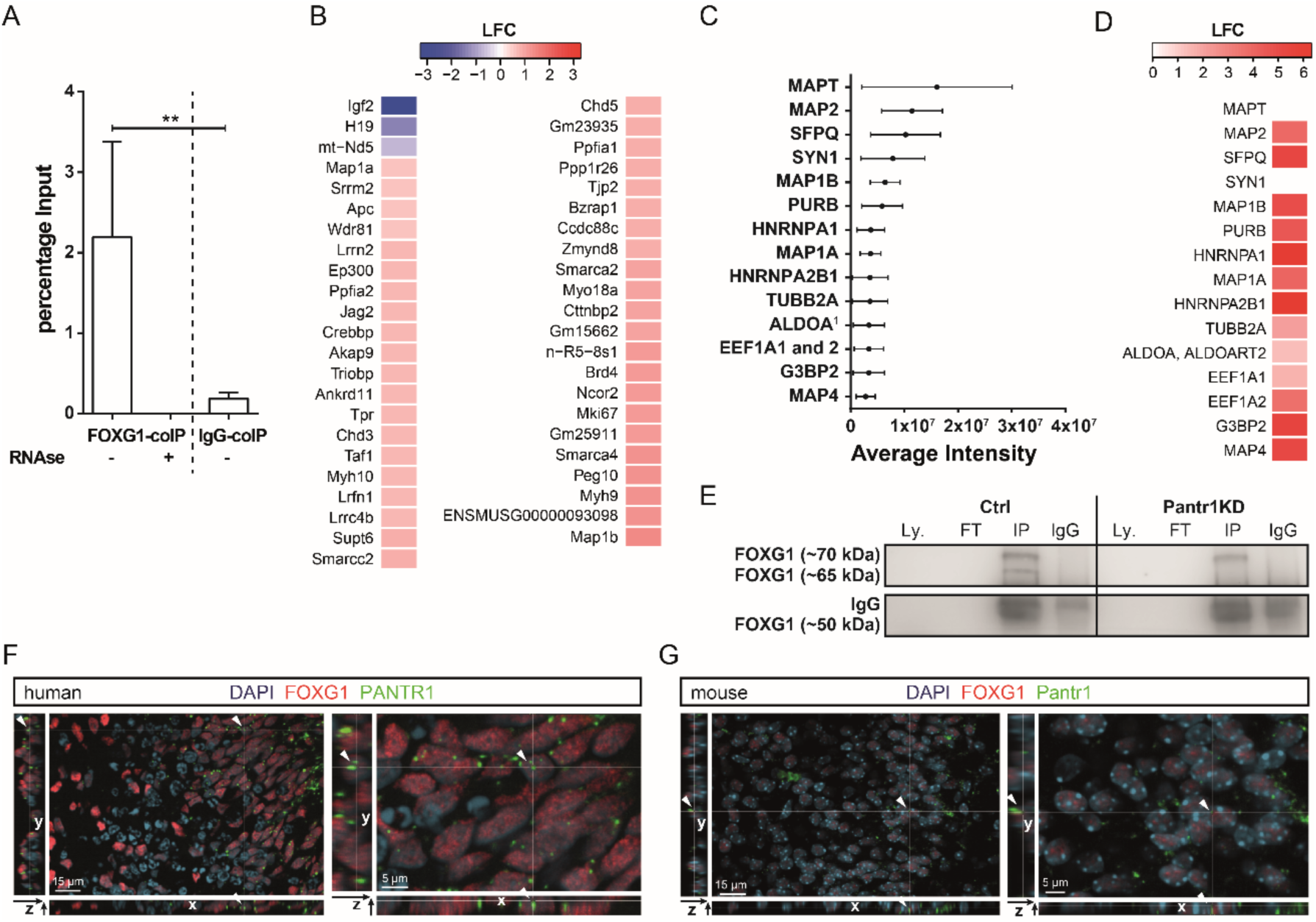
FOXG1 associates with *Pantr1*. **(A)** RIP-qRT-PCR. Shown is the percentage of the input for *Pantr1* after co-precipitation with FOXG1 or IgG in presence or absence of RNAse of whole brain lysates of P0 wildtype mice. n=3, ratio paired Student’s t-test: *: p < 0.05. **(B)** Enriched genes after RNA immunoprecipitation sequencing (RIPseq) using anti-FOXG1 coated beads compared to IgG coated beads. Reads were mapped against murine genome (mm10). The counts of features were normalised and statistically analysed with Deseq2. Shown are the genes with adjusted p-value (padj) < 0.05 and a LFC < -0.584 or > 0.584. n=3. **(C)** Proteins with the highest average intensity after MS analysis of *in vitro* transcribed *Pantr1* RNA pull down using 6-week adult mouse hippocampus protein lysates. Only proteins with at least 2 identified peptides, a coverage over 10% and a p-value below 0.05 were taken into account. Given is the average intensity with SEM. n=3. **(D)** Heatmap of the FOXG1 interactome enrichment of proteins shown to bind to *Pantr1* in the RNA pull down assay. Full dataset of the FOXG1 interactome is published in (23). **(E)** Immunoblot after immunoprecipitation of FOXG1 from the lysate of primary hippocampal neurons (DIV7) treated either with shCtrl (Ctrl) or ShPantr1_1 (*Pantr1*KD). Shown are the different FOXG1 bands occurring in the cell lysate (Ly.), the supernatant of the antibody conjugated beads (Flow through, FT), the proteins bound to the FOXG1 antibody conjugated beads (IP) and the proteins bound to the IgG antibody conjugated beads as negative control (IgG). n=3. **(F)** hCO stained with smFISH probe targeting *PANTR1* (green) and immunostained for FOXG1 (red). The image shows the orthogonal projections of confocal z-stacks. The orientation of the x-, y- and z-axis is indicated. The arrowheads indicate an example of colocalization of *PANTR1* and FOXG1. **(G)** Mouse E18.5 brain slide stained with smFISH probe targeting *Pantr1* (green) and immunostained for FOXG1 (red). The image shows the orthogonal projections of confocal z-stacks. The orientation of the x-, y- and z-axis is indicated. The arrowheads indicate an example of colocalization of *Pantr1* and FOXG1. On the right side, a magnification of the left image is shown.

### FOXG1 and *Pantr1* synergise on transcriptional regulation of genes involved in neuronal differentiation

Combining our results so far, we hypothesised that FOXG1 and *Pantr1* might converge on transcriptional regulation. To explore altered transcriptomes, we transduced mouse DIV (days *in vitro*) 7 primary hippocampal neurons with shRNAs to knockdown (KD) either *Foxg1, Pantr1,* or luciferase control. We confirmed the KD by qRT-PCR (**Fig. 5A**). KD of *Foxg1* let to 2971 differentially expressed genes (DEGs), of which 1203 increased and 1768 decreased compared to the control, as determined by RNAseq and using a threshold of +/- 0.58 log_2_(foldchange) (LFC) and padj ≤ 0.05. For shPantr1, we identified in total 1800 DEGs (1254 decreased, 546 increased) compared to control (**Fig. 5B**). Whereas *Pantr1* was among the decreased DEGs (LFC= -0.435, padj= 0.020) upon *Foxg1*KD, *Foxg1* was not among the DEGs upon *Pantr1*KD (LFC= 0.036, padj= 0.755), confirming FOXG1 as an upstream regulator of *Pantr1*. In total, we detected 2149 genes unique to *Foxg1*KD, 978 to *Pantr1*KD, and 822 genes were affected in common in both KD conditions (**Fig. 5C, D**). With the 822 common DEGs, 232 increased and 532 decreased in both KD conditions, whereas 47 increased upon *Foxg1*KD, but decreased upon *Pantr1* reduction, and 11 decreased in *Foxg1*KD, but increased upon *Pantr1*KD. GO-term enrichment analysis showed that decreasing DEGs upon *Foxg1*KD or *Pantr1*KD were functionally involved in neuronal differentiation (**Fig. 5E**). As suspected, the transcriptional changes upon *Foxg1*KD or *Pantr1*KD indicated a common regulative network impacting neuronal differentiation in the mouse hippocampus.

**Fig. 5:**
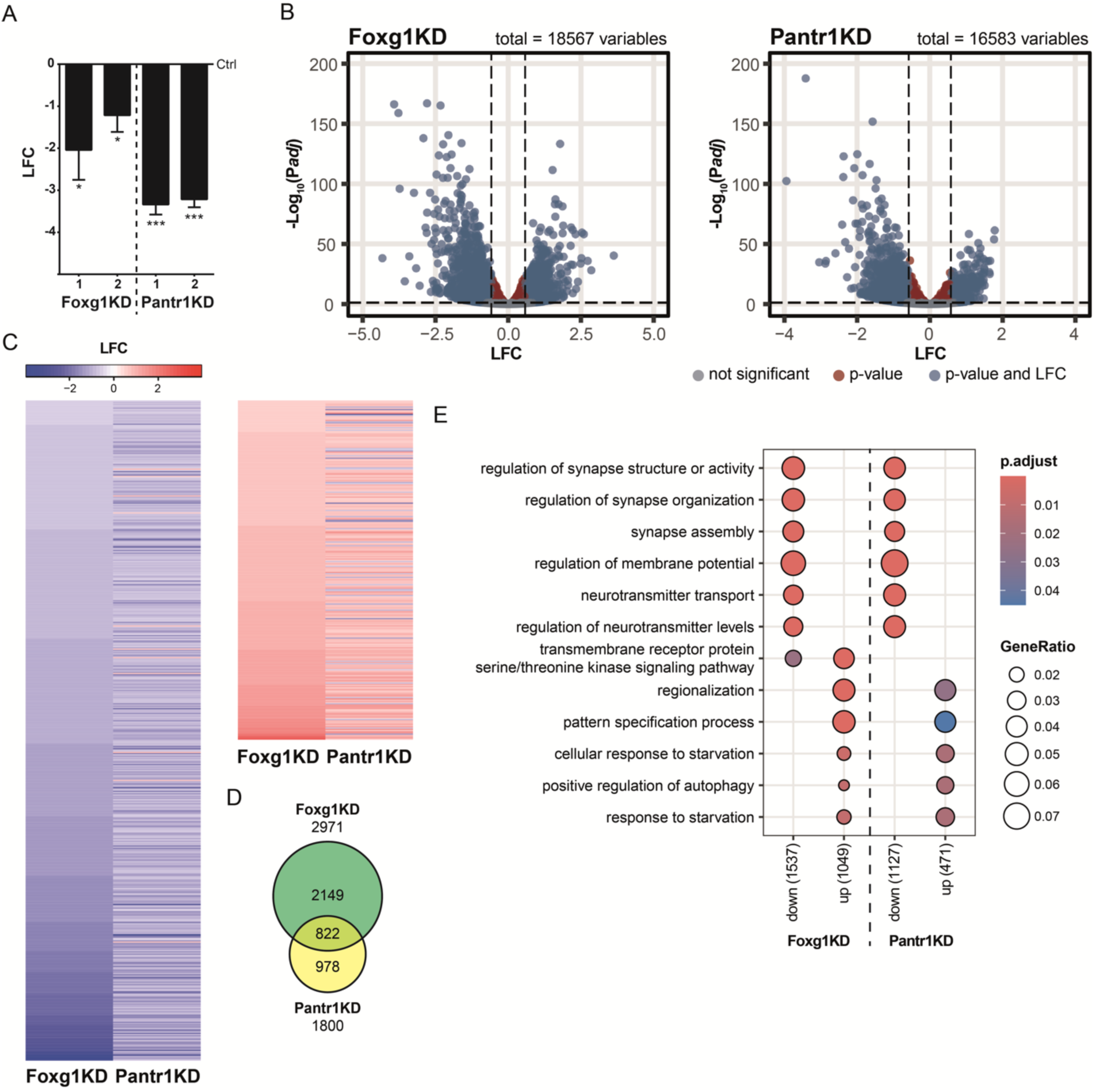
FOXG1 and *Pantr1* share transcriptional target genes impacting neuronal differentiation. **(A)** Primary hippocampal neurons DIV7 transduced with either shCtrl or shRNAs targeting *Foxg1* or *Pantr1*. All transcript levels decreased significantly upon KD with the respective shRNAs (shFoxg1_1, shFoxg1_2; and shPantr1_1, shPantr1_2) compared to shCtrl, as assessed by qRT-PCR with *Gapdh* as housekeeping gene. Given is the log_2_(foldchange) (LFC) of the expression in shRNA treated cells (*Pantr1*KD and *Foxg1*KD) compared to shCtrl treated cells (Ctrl). n=3 (shFoxg1_1 and shPantr1_1), n=4 (shFoxg1_2 and shPantr1_2). Unpaired Student’s t-test to Ctrl: * p<0.05, ** p<0.01, *** p<0.001. **(B)** Volcano plots of differentially expressed genes (DEG) in DIV7 primary hippocampal neurons transduced with shFoxg1_1 (*Foxg1*KD) compared to shCtrl (Ctrl), and shPantr1_1 (*Pantr1*KD) compared to shCtrl (Ctrl). The log_2_(foldchange) (LFC) compared to the control is shown on the x-axis while the y-axis is showing the mean expression value of -log10 (p-value). Genes with increased or decreased expression and an adjusted p-value (padj) < 0.05 (red) and additionally LFC < -0.584 or > 0.584 (blue) based on the Deseq2 analysis results are indicated. n=5. Dashed lines are indicating the thresholds for padj (0.05) and LFC (−0.584 and 0.584). **(C)** Heatmap of DEGs upon *Foxg1*KD and *Pantr1*KD in primary hippocampal neurons. Given is the log_2_(foldchange) (LFC) for common DEGs with an adjusted p-value (padj) < 0.05 and a LFC < -0.584 or > 0.584. n=5. **(D)** Venn diagram of DEGs upon KD of *Foxg1* or *Pantr1* with an adjusted p-value (padj) < 0.05 and a LFC < -0.584 or > 0.584, indicating a set of 822 commonly regulated target genes. n=5. **(E)** GO-term enrichment analysis of DEGs upon KD of *Foxg1* or *Pantr1* split according to DEGs with either increased or decreased expression compared to shCtrl. Given are the top enriched GO terms with a FDR > 0.5.

### *Pantr1* localises FOXG1 at specific binding sites at the chromatin level

Since (i) RIP-qRT-PCR and RIPseq showed association of FOXG1 and *Pantr1*, (ii) *Pantr1* was involved in forming FOXG1^65^, and (iii) FOXG1 and *Pantr1* had common transcriptional targets, we hypothesised that *Pantr1* could potentially affect the localisation of FOXG1 at the chromatin level. To test this hypothesis, we performed ChIPseq of FOXG1 upon *Pantr1*KD in primary hippocampal neurons and identified indeed significantly differentially bound regions (DBRs) of FOXG1 compared to controls (**Fig. 6A** for significant peaks; all called FOXG1 peaks displayed in **Fig. S4A**). Among the significant DBRs 233 increased and 21 decreased FOXG1 binding upon *Pantr1* reduction. To the largest part, DBRs had promoter or distal intergenic features (**Fig. 6B**). GO-term enrichment retrieved functional terms associated with neuronal differentiation (**Fig. 6C**; all called FOXG1 peaks displayed in **S4B**), in accordance with the observed transcriptional regulations upon reduced expression of FOXG1. We identified 13 shared DEGs that also harboured significant DBRs (**Fig. S5**), which rendered these as *Pantr1*-dependent FOXG1 target genes (**Fig. 6D**; common DEGs after *Foxg1*KD and DBRs upon *Pantr1*KD displayed in **Fig. S4C**). Expression of these *Pantr1*-dependent FOXG1 targets increased or decreased, and this bimodal impact of FOXG1 on transcription was in accordance with our previous observations (22). We also observed enrichment of FOXG1 in human d105 hCO in most of the 13 mouse ChIPseq target genes. In part the FOXG1 enrichment was found at sites for which we identified homology between human and mice, and which were differentially bound by FOXG1 upon *Pantr1*KD in mouse cells (**Fig. S6**). It is thus tempting to speculate that *PANTR1* is involved in site-specific FOXG1 localisation in humans in similar manner as observed in the mouse. Supporting this hypothesis is the finding that *SLC8A1*, an important mediator of intracellular sodium/calcium homeostasis (45), decreased in both human and mouse models with reduced levels of FOXG1 (**Fig. S6**).

**Fig. 6:**
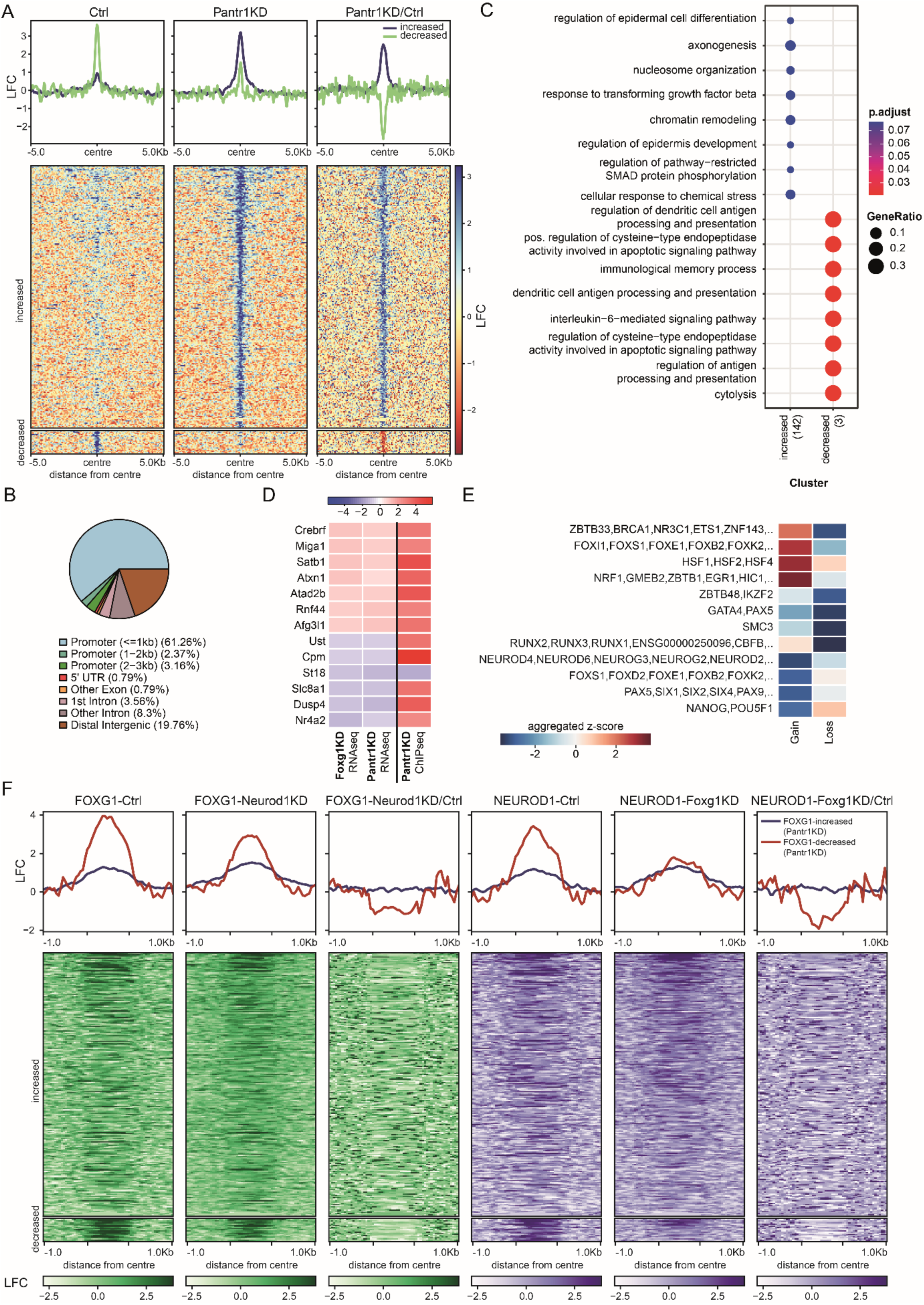
*Pantr1* regulates FOXG1 presence at the chromatin. **(A)** Heatmap of differently bound regions (DBRs) of FOXG1 upon *Pantr1*KD, 5 Kb up-/downstream of FOXG1 peak summits in control and *Pantr1*KD conditions. DBRs were calculated using Diffbind. Data are normalised by sequencing depth and input control as log_2_(ChIP/Input) for control and *Pantr1*KD data. The difference between *Pantr1*KD and control conditions was calculated from RPKM normalised bigwig files as log_2_ (*Foxg1*KD/Control). The metaprofiles (top) show the mean log_2_(foldchange) (LFC) of increased (blue) and decreased (green) regions. n=2. **(B)** Genome-wide distribution of DBRs of FOXG1 upon *Pantr1*KD in DIV7 primary hippocampal neurons in percentage for near and far promotor regions, 5’UTR, intronic and exonic as well as distal intergenic regions. **(C)** GO-term enrichment of differentially bound regions of FOXG1 upon *Pantr1*KD in DIV7 primary hippocampal neurons with a qvalue cutoff of 0.05 and pvalue cutoff of 0.05. **(D)** Heatmap of common DEGs in the RNAseq data sets after *Pantr1*KD and *Foxg1*KD (adjusted p-value (padj) < 0.05 and a LFC < -0.584 or > 0.584) and annotated genes (ChIPSeeker) corresponding to the significant DBR in the FOXG1-ChIPseq data set after *Pantr1*KD. Shown are the LFC (RNAseq) or fold change (ChIPseq) compared to the control condition. **(E)** Motifs analysis of the increased (gain) and decreased (loss) DBRs. Shown is the aggregated z-score. **(F)** Heatmap of FOXG1 (green) and NEUROD1 (purple) enrichments at the increased and decreased DBRs under control, *Neurod1*KD, and *Foxg1*KD conditions. Data were obtained from (22) and were normalised by sequencing depth and input control as log2(ChIP/Input). The difference between *Foxg1*KD-Control and *Neurod1*KD-Control conditions were calculated from RPKM normalised bigwig files as log2(*transcription factor* KD/Control). The metaprofiles (top) show the average log_2_(foldchange) (LFC) of increased (blue) and decreased (red) regions. n=2.

We performed a differential transcription factor DNA-binding motif enrichment analysis for the significant FOXG1 DBRs upon *Pantr1*KD in mouse hippocampal cells. Increased binding of FOXG1 occurred at sites that were enriched in GC- and GAA-rich motifs, characteristic for Zinc-finger- or heat shock transcription factors, in addition to generic Fkh-binding motifs encompassing the entire Fkh family (**Fig. 6E**). On the other hand, regions with NEUROD-(bHLH) binding sites or palindromic Fkh motifs, bound specifically by FOXO and FOXA family members, were fewer among the motifs under bound sites (**Fig. 6E, S4D** for all called peaks). These findings suggested that upon *Pantr1*KD, FOXG1 was redistributed at the chromatin, seemingly favouring generic sites harbouring a variety of motifs.

Motifs conferring binding sites of transcription factors known to interact with FOXG1, including FOXO members and NEUROD1, were comparably less represented in the regions gaining FOXG1 binding upon *Pantr1*KD. We recently reported on the cooperative presence of FOXG1 and NEUROD1 at several FOXG1 bound regions (22), and observed decreased occurrence of FOXG1 binding at NEUROD motifs upon *Pantr1*KD. Therefore, we hypothesised that regions where FOXG1 binds in a *Pantr1*-dependent manner also exhibited a specific pattern of NEUROD1 binding. To test this hypothesis, we specifically analysed *Pantr1*-dependent FOXG1 bound regions in the NEUROD1 ChIPseq dataset. At DBRs with increased binding of FOXG1 upon *Pantr1*KD, we did not observe significant alterations in the binding pattern of FOXG1 or NEUROD1 upon KD of either *Foxg1* or *Neurod1*. However, focusing on regions where FOXG1 binding decreased upon *Pantr1*KD, we observed a corresponding reduction in NEUROD1 binding upon *Foxg1*KD and a decrease in FOXG1 binding upon *Neurod1*KD (**Fig. 6F**). Together, these findings suggested that *Pantr1* facilitates cooperative presence of FOXG1/NEUROD1 at specific sites, and *Pantr1* reduction leads to redistribution of FOXG1 to comparably more generic binding sites.

### *Pantr1* OE upon reduced expression of *Foxg1* rescues dendritic outgrowth defects

Since GO-term enrichment suggested that impaired neuronal differentiation might be a common feature of reduced expression of *Foxg1* or *Pantr1,* we transduced primary hippocampal neurons with the respective shRNAs, cultured them until DIV7 and quantified dendritic branching and outgrowth using Sholl-analysis. The number of dendrites decreased 10 to 100 µm proximal to the soma upon either *Foxg1*KD or *Pantr1*KD condition compared to control (**Fig. 7A, B**). Further, we tested whether *Pantr1*KD would impair spine density and spontaneous excitatory post-synaptic currents (sEPSC), as reported for *Foxg1* loss-of-function (16). Indeed, we observed a reduced spine density in transduced primary hippocampal cells that expressed shPantr1 alongside a GFP reporter construct compared to control condition (**Fig. 7C, D**). This decrease in spine density was associated with reduced spontaneous firing rates (**Fig. 7E**), decreased sEPSC amplitudes (**Fig. 7F**) and a lower frequency of sEPSCs (**Fig. 7G**) following *Pantr1*KD compared to controls. These findings are consistent with a downregulation of *Gria2* (Glutamate Ionotropic Receptor AMPA Type Subunit 2) and *Gria4* expression upon *Pantr1*KD as indicated by the transcriptome data (**Fig. 7H**).

**Fig. 7:**
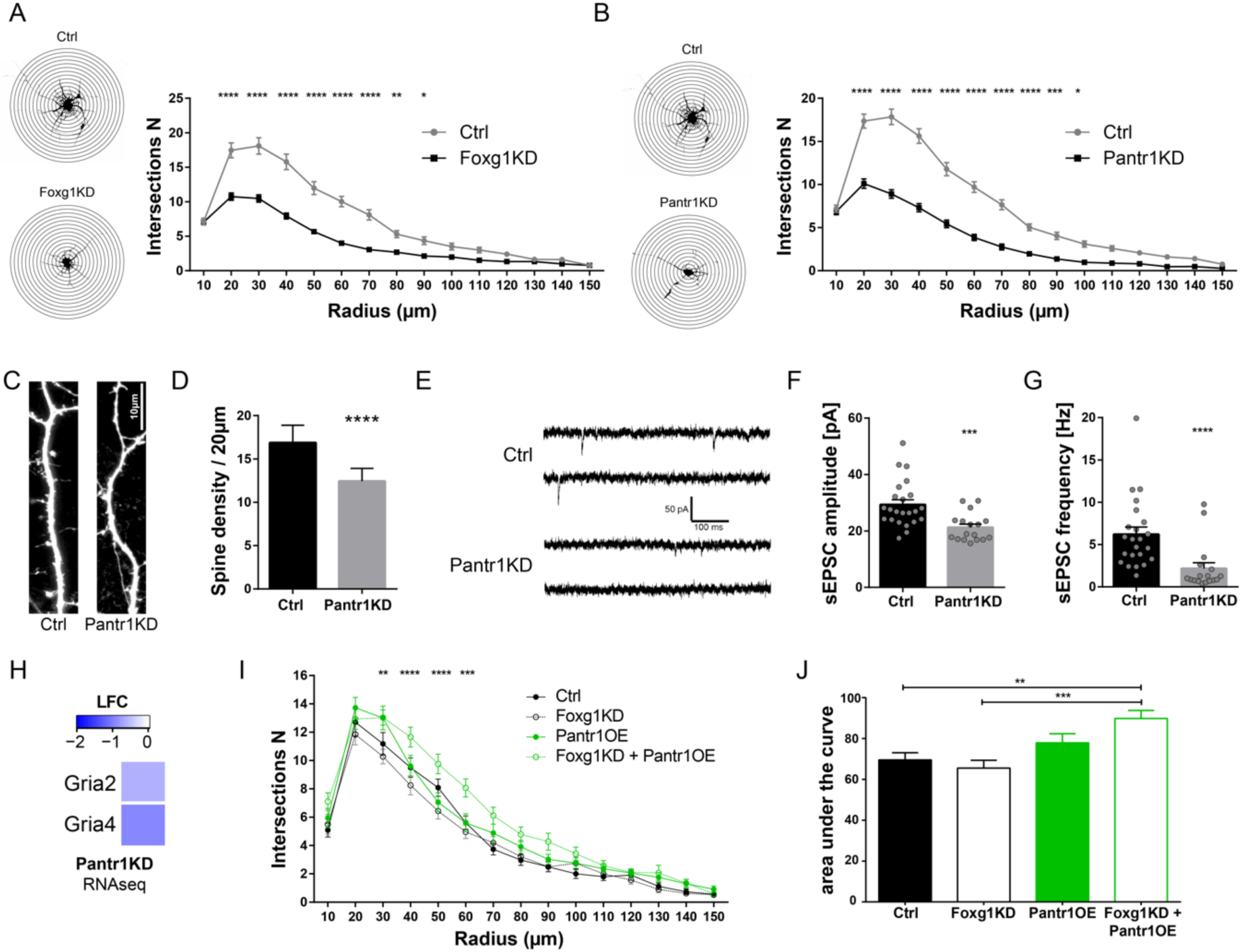
*Pantr1* partially rescues neuronal differentiation defects upon FOXG1 reduction. **(A, B)** Camera lucida reconstruction of representative DIV7 primary neurons after transduction with shCtrl (Ctrl), shFoxg1 (Foxg1KD) or shPantr1 (Pantr1KD) and superimposed concentric circles used for Sholl-analysis. The radius interval between the circles was 10 µm per step. Quantification of Sholl-analysis of control (shCtrl, grey) and *Foxg1*KD (**A**, combined for shFoxg1_1 and 2; black) or *Pantr1*KD (**B**, combined for shPantr1_1 and 2; black), showing significantly decreased dendritic complexity upon *Foxg1*KD (shCtrl: n=45, shFoxg1_1 and 2: n=76) or *Pantr1*KD (shCtrl: n=65, shPantr1_1 and shPantr1_2: n=53). Two-way ANOVA, Sidak multiple comparison test. *: p < 0.05, **: p < 0.01, ***: p < 0.001, ****: p < 0.0001. **(C)** Representative images of spines from dendritic branches upon shCtrl and shPantr1_1. Biocytin-filled neurons were stained with Alexa Fluor™ 633-conjugate. Scale bar: 10 μm. **(D)** Quantification of spines in 20 µm increments of stained dendrites as shown in C, showing that spine density reduced significantly in *Pantr1*KD condition (shCtrl: 13 cells from 4 cultures; shPantr1_1: 19 cells from 4 cultures, unpaired Student’s t-test, ****: p < 0.0001). **(E)** Representative traces of spontaneous excitatory postsynaptic currents (sEPSCs) from shCtrl or shPantr1_1 transduced hippocampal neurons recorded on DIV14. **(F,G)** Group data of mean sEPSC amplitude (**F**) and frequency (**G**) from *Ctrl* and *Pantr1*KD neurons recorded on DIV14. shCtrl: n=23, shPantr1_1: n=17. Significance was determined by Mann-Whitney test, ***p < 0.001, ****p < 0.0001 (H) Heatmap of log_2_(foldchange) (LFC) expression level changes of *Gria2* and *Gria4* upon *Pantr1*KD in DIV7 hippocampal neurons compared to controls as revealed by RNAseq analysis. (I) Sholl analysis of dendrite complexity upon *Foxg1*KD (black, dashed line), *Pantr1* overexpression (OE) (green line) and combination of *Foxg1*KD and *Pantr1*OE (green, dashed line) compared to Ctrl (black line), showing that *Pantr1*OE rescues reduced dendritic complexity upon FOXG1 reduction (Ctrl n=34, *Foxg1*KD n=32, Pantr1OE n=33, *Foxg1*KD_*Pantr1*OE n=29). The radius interval between the circles was 10 µm per step. Two-way ANOVA with Sidak multiple comparison test was performed over all conditions. Shown is the significance of *Foxg1*KD_*Pantr1*OE vs *Foxg1*KD, **p < 0.01, ***p < 0.001, ****p < 0.0001. (J) Bar chart representation of quantifications given in **I**. Given is the sum of intersections of all intervals as area under the curve for control cells and cells with *Foxg1*KD, *Pantr1* overexpression (OE) and the combination of *Foxg1*KD and *Pantr1*OE. One-way ANOVA with Tukey’s multiple comparison test was performed over all conditions, **p < 0.01, ***p < 0.001.

Our data suggested that reduced expression of FOXG1 decreased *Pantr1* expression, and that the regulative network centred on both factors impacted dendritic outgrowth in hippocampal neurons. The upstream action of FOXG1 implied that overexpression of *Pantr1* would be a means to rescue - at least in part - loss of FOXG1 function. To address this potential functional interplay between FOXG1 and *Pantr1*, we expressed either shFoxg1 alone, overexpressed *Pantr1* or expressed shFoxg1 alongside *Pantr1* overexpression (OE) in primary hippocampal neurons. Sholl-analysis of hippocampal neurons showed a mild, but still observable decrease in dendritic arborisation upon *Foxg1*KD, but significant increase upon *Pantr1*OE in the condition of *Foxg1*KD, compared to the control (**Fig. 7I**). The rescue effect was strongest in a range near the soma, between 10 and 60 μm (**Fig. 7I, J**). This data suggested that in the context of dendritic development *Pantr1* acts downstream of FOXG1 to regulate this aspect of neuronal differentiation in mouse hippocampal neurons.

## DISCUSSION

Our investigation into the role of l(i)ncRNAs in the context of FOXG1-syndrome led to identifying *Pantr1* as one of the crucial lincRNAs that shows altered expression levels across various model systems exhibiting *Foxg1*-haploinsufficiency or full deletion. Here, we report a functional synergy between FOXG1 and *Pantr1* that in our study presents with two distinct molecular mechanisms. For one, our transcriptomic and functional characterisation place FOXG1 as an upstream regulator of *Pantr1* expression in the cerebral cortex and hippocampus of the mouse as well as in hCO. In hCO, FOXG1 deletion impaired proper *PANTR1* expression especially in distinct interneuron stages. It might be tempting to speculate that dysregulated *PANTR1* levels correlate with the disbalance of excitatory and inhibitory activity in iPSC-derived model systems of human FOXG1-syndrome patients (2,17). Notably in the context of the human FOXG1-syndrome is that the overexpression of *Pantr1* in the condition of reduced FOXG1 levels in mouse hippocampal neurons rescued dendritic complexity. This finding renders the lincRNA *Pantr1* as an important mediator of FOXG1 function in promoting the correct formation of the dendritic tree in the mouse; but it might also be a potential target for testing therapeutic options in humans. However, much more research on *PANTR1* functions especially in the human system in the different cell subtypes is needed in further upcoming studies.

The second molecular mechanism upon which FOXG1 and *Pantr1* converge results from our observation that FOXG1 is associated with *Pantr1*. Together with the recent report that FOXG1 associates with L1 mRNA (44), it seems emerging that FOXG1 experiences both DNA- as well as RNA-associations, despite being devoid of canonical RNA-binding motifs. Recent reports link RNA binding to other features than classical RNA-binding domains, including intrinsically disordered regions (IDR), and they suggest interaction of DNA/DNA-binding proteins with RNA-binding proteins (46-48). Along these lines, the fkh protein domain has been recognised mainly as DNA-binding so far, but recently it has been also reported to confer association with RNA by binding to G2-rich sequences, being composed of a consensus sequence containing 8 GG nucleotides interspersed by one or two different nucleotides (43). We did not identify a G2-rich consensus within the sequences of human *PANTR1* (ENST00000443988.7, PANTR1-202) or mouse *Pantr1* (ENSMUSG00000060424, all exons). Nevertheless, this observation does not exclude direct binding between FOXG1 and *Pantr1*, because the RNA-binding capability of FOXG1 might be encoded in its extensive IDR, as predicted by using disoFLAG (49). However, the lack of consensus might also support the hypothesis of another protein interactor that associates with both FOXG1 and *Pantr1*, facilitating their interaction.

In support of this assumption, our FOXG1 interactome and RIP followed by mass spectrometry revealed that the common protein interaction partners shared between FOXG1 and *Pantr1* are mainly described as being cytoplasmic proteins and contain several RNA-binding proteins. Our study showed that presence of *Pantr1* coincided with a specific FOXG1 protein isoform, with a slightly higher molecular mass, i.e. FOXG1^65^, compared to slightly lower, predicted mass of FOXG1 in FOXG1^50^. Association with a small, 10-15 kDa protein could explain this shift in mass. However, neither the interactome nor RIP followed by mass spectrometry studies suggested a common top enriched protein with 10-15 kDa that could account for the difference between FOXG1^50^ and FOXG1^65^. It thus remains to be explored whether FOXG1^65^ contains RNA, represents a stable conformation, or has a small, but currently undetected protein interactor.

The enrichment of cytoplasmic binding partners to FOXG1 and/or *Pantr1* could result from a technical bias, for example caused by predominant extraction of cytoplasmic proteins for the *Pantr1* binding assay. On the other hand, predominant cytoplasmic interactors can reflect that FOXG1 would be sequestered in the cytoplasm through *Pantr1*. A sequestering function of *Pantr1* for FOXG1 could explain the increased binding of FOXG1 to chromatin upon *Pantr1*KD. However, the sponging being the sole purpose of FOXG1-*Pantr1* complexes is not well supported by two of our observations: (i) the nuclear colocalisation of FOXG1 and *Pantr1 in vivo*, and (ii) *Pantr1*’s sponging effect on FOXG1 protein in the cytoplasm would scarcely account for the selectivity observed in differentially bound chromatin regions of FOXG1 upon *Pantr1*KD.

The limited number of loci, with mainly increased FOXG1 binding, rather suggests that *Pantr1* is involved in localising FOXG1 by conferring site-specific transcription factor binding. In this context, our data suggest that *Pantr1* does not interfere with *de novo* binding of FOXG1 to the chromatin, as sites with increased binding had FOXG1 peaks also in the control condition. Rather our data indicate that *Pantr1* limits (i) FOXG1 presence at already bound regions (**Fig. S4**) and (ii) spreading of FOXG1 to generic binding sites (**Fig. 6E**). On the other hand, *Pantr1* might facilitate cooperative presence of FOXG1 and NEUROD1, as the regions where FOXG1 binding decreased upon *Pantr1*KD coincide with regions where loss of either FOXG1 or NEUROD1 affects the localisation of the other transcription factor (**Fig. 6F**).

In conclusion, based on our data, we propose that *Pantr1* modulates FOXG1 presence at the chromatin. *Pantr1* functions to destabilise FOXG1 binding and to prevent its spreading into generic chromatin regions, while also facilitating FOXG1 binding at sites of cooperative NEUROD1 binding.

Clearly, much more research is needed to unravel how FOXG1 localisation within cellular compartments, at the chromatin level, and in association with RNAs is regulated. Evidently, our study of *Pantr1* localisation and interaction with putative other protein partners suggest a plethora of additional functions for *Pantr1* as well as FOXG1 in the cytoplasm and nucleus that await further research.

## DATA AVAILABILITY

RNAseq data after scRNAseq of cortical organoids (CO) from healthy donor (HD) and FOXG1del were deposited to the NCBI Gene Expression Omnibus (GEO) with accession number GSE273180. Human FOXG1-ChIPseq from hCO were deposited to GEO with the accession number GSE272853. RNAseq data of E18.5 hippocampus and adult hippocampus were published previously (22,23), and have the accession numbers GSE189119 (E18.5 hippocampus) and GSE106801 (adult hippocampus). E13.5 RNAseq data from cerebral cortex have the accession number GSE95833. RNAseq data from DIV7 primary hippocampal neurons with Foxg1KD and Pantr1KD were deposited to GEO with accession number GSE172220. Murine FOXG1-ChIPseq data after Pantr1KD in DIV7 primary hippocampal neurons were deposited to GEO with accession number GSE272852. RIPseq data were deposited to GEO with accession number GSE272854. NEUROD1 and FOXG1-ChIPseq data of DIV11 primary hippocampal neurons are published with accession number GSE189119 (22). All other data types and codes recreating the analyses from the data files can be found at DOI: 10.6084/m9.figshare.26788915 (https://figshare.com/s/bbff462c5689515bfb39) as R markdown files, Python scripts, and Galaxy workflows.

## Tokens for reviewers

GSE272852

GSE272853

GSE272854

GSE273180

GSE172220

## SUPPLEMENTARY DATA

Supplementary Data are available online.

## AUTHOR CONTRIBUTIONS

TV conceived and designed the study. Material preparation, data collection, and analysis were performed by FG, TR, IA, CLF, GA, and TV. The first draft of the manuscript was written by TV together with FG and IA, and all authors read and approved the final manuscript.

## ACKNOWLEDGEMENT

The authors thank S. Heidrich for technical assistance. We also thank A. Izzo for scientific support and critical comments on the manuscript. We thank E. Gabriel and J. Gopalakrishnan, Heinrich-Heine-University Düsseldorf, Germany, for scientific support in the generation of organoids. The authors thank M. Börries, Institute of Medical Bioinformatics and Systems Medicine, University Medicine Freiburg, for infrastructural support of the study. The authors acknowledge the support of the Freiburg Galaxy Team: P. Videm and B. Grüning, Bioinformatics, University of Freiburg, Germany, funded by the German Federal Ministry of Education and Research BMBF grant 031 A538A de.NBI-RBC and the Ministry of Science, Research and the Arts Baden-Württemberg (MWK) within the framework of LIBIS/de.NBI Freiburg. We also thank U. Bönisch, MPI Immunobiology and Epigenetics, Freiburg, Germany, from the sequencing core facility. Control hiPSCs were provided by the FiPS Core Facility, University Medicine Freiburg, Germany. The “Cell lines and DNA bank of Rett Syndrome, X-linked mental retardation and other genetic diseases”, member of the Telethon Network of Genetic Biobanks (project no. GTB12001), funded by Telethon Italy and of the EuroBioBank network, provided us with hiPSC with FOXG1 deletion.

## ETHICS APPROVAL

All mouse experiments were approved by the animal welfare committees of the respective authorities (X-14/04H, X-17/03S and X-22/02B, Regierungspräsidium Freiburg, Germany).

## SUPPLEMENTARY FIGURE LEGENDS

**Fig. S1: Characterisation of human brain organoids and expression of FOXG1**

**(A)** Immunostainings of control organoid slices after 105 days in culture presenting different markers (Paired box protein (PAX6), Neuron-specific class III beta-tubulin (TUJ), neuronal nuclei (NeuN), glial fibrillary acidic protein (GFAP), vesicular glutamate transporter 1 (vGLUT1), vGLUT2, T-box brain transcription factor 1 (TBR1), TBR2) of neural development. Scale bars as indicated in the figures.

**(B)** Immunostainings of FOXG1 in CO of healthy donor (HD) and FOXG1del showing reduced FOXG1 protein expression in FOXG1del compared to HD after 105 d ays in culture. Dashed line delineates the organoid surface. Scale bars as indicated in the figure.

**Fig. S2: FOXG1 associates with RNA**

**(A)** Upper part: Different transcript variants of *Map1b* (orange). Boxes represent the different exons.

Lower part: RIPseq tracks of log_2_(foldchange) (LFC) of reads in FOXG1-coIP vs IgG-coIP in the different n (green) and the average of those LFCs (black) in the genomic region of *Map1b.* On the x-axis the position on the corresponding chromosome is indicated in bp of the mm10 genome.

**(B)** Upper part: Different transcript variants of *Pantr1* (orange). Boxes represent the different exons.

Lower part: RIPseq tracks of log_2_(foldchange) (LFC) of reads in FOXG1-coIP vs IgG-coIP in the different n (green) and the average of those LFCs (black) in the genomic region of *Pantr1.* On the x-axis the position on the corresponding chromosome is indicated in bp of the mm10 genome.

**Fig. S3: FOXG1 localises in the vicinity of *PANTR1* in vivo**

**(A)** Overview of the expression of *PANTR1* (smFISH) and FOXG1 protein in human d105 brain organoids. The boxed area indicates the magnification shown in Fig. 4F.

**(B)** Mouse brain (P0) stained with smFISH probe targeting *Pantr1* (green) and immunostained for FOXG1 (red). Boxed area indicates the magnification shown in Fig. 4G. Scale bars as indicated in the figure.

**Fig. S4: Analyses of FOXG1-ChIPseq peaks upon *Pantr1*KD in DIV7 primary mouse hippocampal neurons**

**(A)** Heatmap of regions 5 Kb up-/downstream of all identified FOXG1 peak summits, represented as k-means clusters with k=4 in control (Ctrl) and *Pantr1*KD condition as well as the comparison of both conditions. Data are normalised by sequencing depth and input control as log_2_(ChIP/Input) for Ctrl and *Pantr1*KD data. The difference between *Pantr1*KD and control condition was calculated from RPKM normalised bigwig files as log_2_ (Pantr1KD/Control). The metaprofiles (top) show the mean log_2_FC (LFC) of each cluster. n=2.

**(B)** GO-term enrichment of the genes in proximity to the FOXG1 ChIPseq summits broken down to the different clusters upon *Pantr1*KD in DIV7 primary hippocampal neurons as shown in (A) with a q-value cutoff of 0.05 and p-value cutoff of 0.05.

**(C)** Heatmap of common DEGs in the RNAseq data sets after *Foxg1*KD (adjusted p-value (padj) < 0.05 and an LFC < -0.584 or > 0.584) and annotated genes (ChIPSeeker) corresponding to all DBRs in the FOXG1-ChIPseq data set upon *Pantr1*KD. Shown are the LFC compared to the control condition. Genes connected to neurite outgrowth and neural development are marked with a yellow box.

**(D)** Motifs analysis of the summits in the different clusters as shown in (A). Shown is a heatmap of the aggregated z-scores.

**Fig. S5: FOXG1 binding profiles at *Pantr1*-dependent FOXG1 target genes in mouse hippocampal cells**

Shown are bigwig tracks of FOXG1 binding sites upon FOXG1 ChIPseq as LFC towards input samples from murine adult hippocampus *in vivo*, and *in vitro* E18.5 cultured primary hippocampal neurons at DIV11 (hipp. primary neurons) (22), E18.5 cultured primary hippocampal neurons at DIV7 (DIV7 control) and upon *Pantr1* KD (DIV7 *Pantr1*KD) as well as the ratio of coverage upon *Pantr1*KD vs control condition (*Pantr1*KD/control). Selected genes classify as *Pantr1*-dependent FOXG1 target genes. The last track displays the binding region called as significant DBR upon *Pantr1* KD. Peak sequences of the DBRs and surrounding loci are marked with a red box. On the x-axis the position on the corresponding chromosome is indicated in bp of the mm10 genome.

**Fig. S6: FOXG1 binding to *Pantr1*-dependent target genes in human cortical organoids**

**(A)** ChIPseq of FOXG1 in human cortical organoids (d105, hCO) showing distribution of enriched FOXG1 peaks of the 13 mouse *Pantr1*-dependent FOXG1 target genes. Red boxes indicate the localisation of the respective DBR of FOXG1 upon *Pantr1*KD in mouse cells as shown in Fig. S6. The last track indicate predicted FOXG1 binding motifs at the respective chromosomal location. On the x-axis the position on the corresponding chromosome is indicated in bp of the hg38 genome. Mapped tracks show the log_2_(foldchange) (LFC) over the input sample for each condition.

**(B)** Violin plots of cell-type specific expression levels of *SLC8A1*. *SLC8A1* expression reduced significantly upon *FOXG1* deletion in ventricular and outer radial glia highlighted in brownish colour, and in the migrating IN II as highlighted in violet.

