## Supplementary information for "The lincRNA *Pantr1* is a FOXG1 target gene conferring site-specific chromatin binding of FOXG1"

**Fig. S6: FOXG1 binding to *Pantr1*-dependent target genes in human cortical organoids**

**(A)** ChIPseq of FOXG1 in human cortical organoids (d105, hCO) showing distribution of enriched FOXG1 peaks of the 13 mouse *Pantr1*-dependent FOXG1 target genes. Red boxes indicate the localisation of the respective DBR of FOXG1 upon *Pantr1*KD in mouse cells as shown in Fig. S6. The last track indicate predicted FOXG1 binding motifs at the respective chromosomal location. On the x-axis the position on the corresponding chromosome is indicated in bp of the hg38 genome. Mapped tracks show the $log_2$(foldchange) (LFC) over the input sample for each condition.

**(B)** Violin plots of cell-type specific expression levels of *SLC8A1*. *SLC8A1* expression reduced significantly upon *FOXG1* deletion in ventricular and outer radial glia highlighted in brownish colour, and in the migrating IN II as highlighted in violet.

figure S1

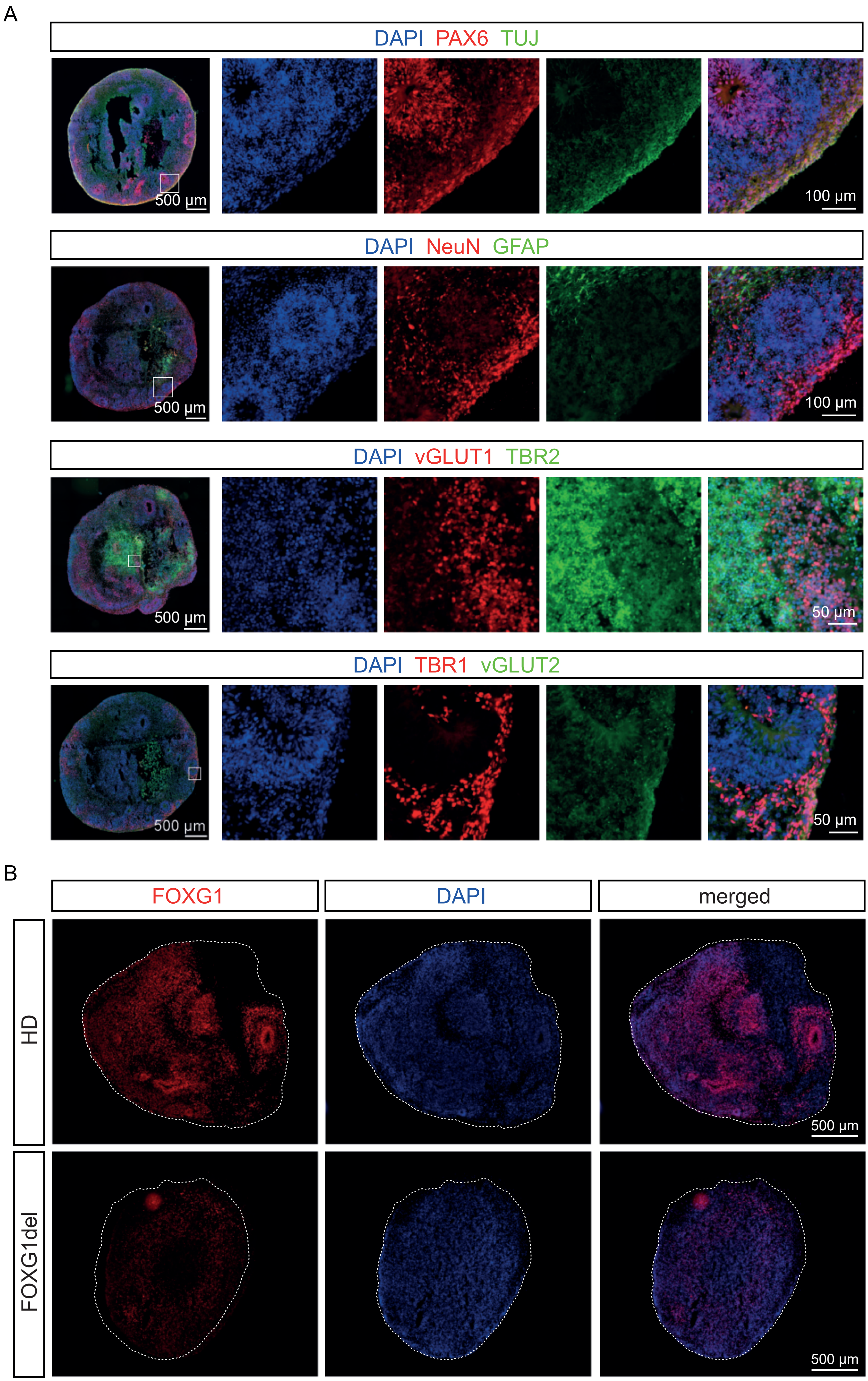

A
DAPI PAX6 TUJ
500 μm
100 μm
DAPI NeuN GFAP
500 μm
100 μm
DAPI vGLUT1 TBR2
500 μm
50 μm
DAPI TBR1 vGLUT2
500 μm
50 μm
B
FOXG1
DAPI
merged
HD
FOXG1del
500 μm
500 μm

figure S2

A

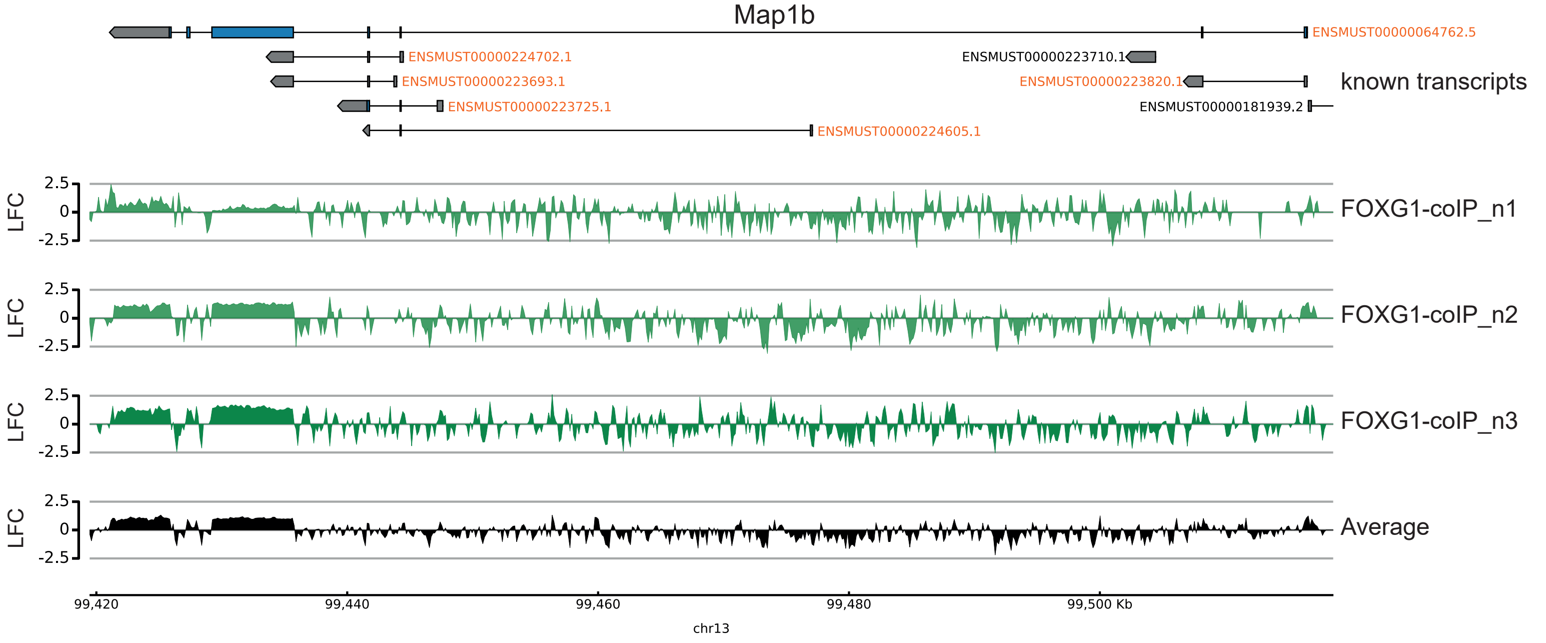
Map1b
ENSMUST00000064762.5
ENSMUST00000224702.1
ENSMUST00000223710.1
ENSMUST00000223693.1
ENSMUST00000223820.1
known transcripts
ENSMUST00000223725.1
ENSMUST00000181939.2
ENSMUST00000224605.1
LFC
2.5
0
-2.5
FOXG1-coIP_n1
FOXG1-coIP_n2
FOXG1-coIP_n3
Average
99,420
99,440
99,460
99,480
99,500 Kb
chr13

B

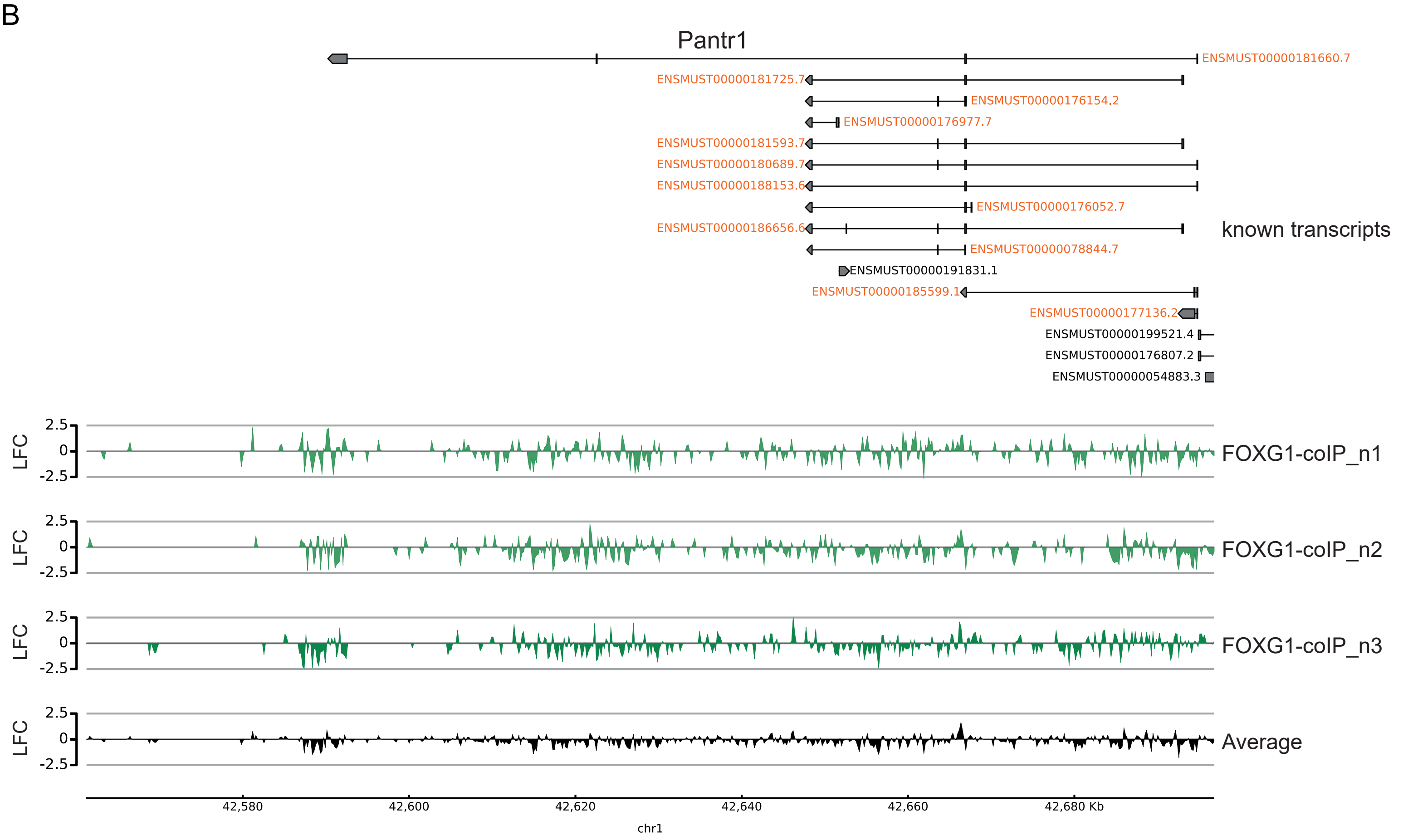
Pantr1
ENSMUST00000181660.7
ENSMUST00000181725.7
ENSMUST00000176154.2
ENSMUST00000176977.7
ENSMUST00000181593.7
ENSMUST00000180689.7
ENSMUST00000188153.6
ENSMUST00000176052.7
ENSMUST00000186656.6
known transcripts
ENSMUST00000078844.7
ENSMUST00000191831.1
ENSMUST00000185599.1
ENSMUST00000177136.2
ENSMUST00000199521.4
ENSMUST00000176807.2
ENSMUST00000054883.3
LFC
2.5
0
-2.5
FOXG1-coIP_n1
FOXG1-coIP_n2
FOXG1-coIP_n3
Average
42,580
42,600
42,620
42,640
42,660
42,680 Kb
chr1

figure S3

A

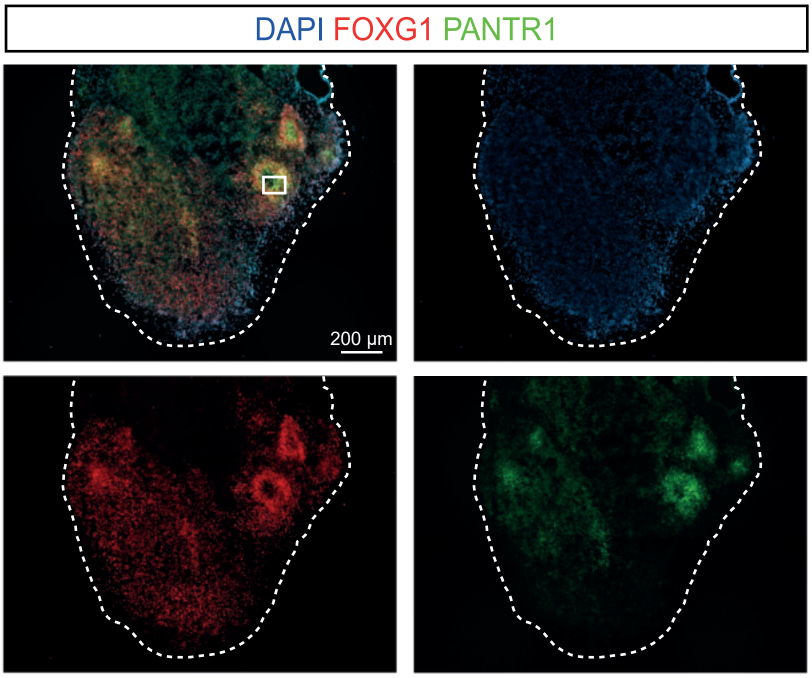


B

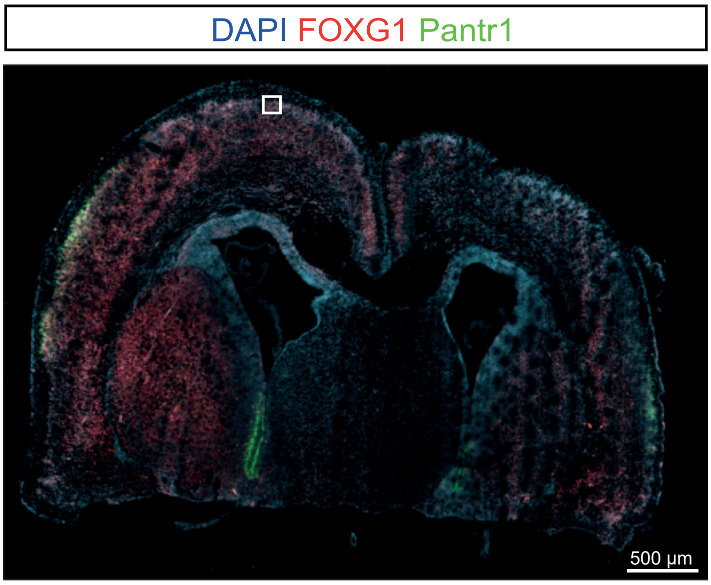

figure S4

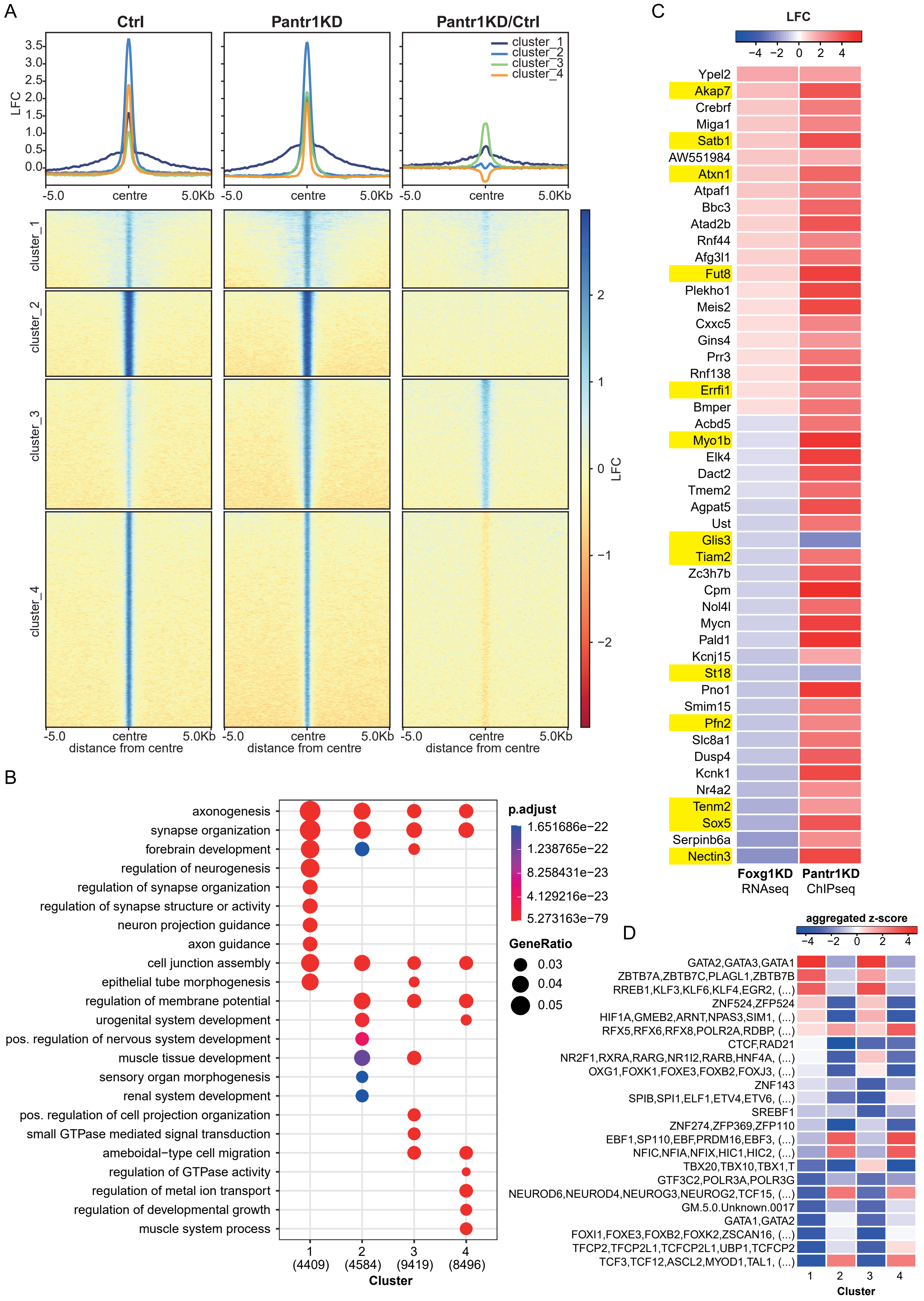
A
Ctrl
Pantr1KD
Pantr1KD/Ctrl
LFC
3.5
3.0
2.5
2.0
1.5
1.0
0.5
0.0
cluster_1
cluster_2
cluster_3
cluster_4
-5.0
centre
5.0Kb
2
1
0
−1
−2
distance from centre
B
axonogenesis
synapse organization
forebrain development
regulation of neurogenesis
regulation of synapse organization
regulation of synapse structure or activity
neuron projection guidance
axon guidance
cell junction assembly
epithelial tube morphogenesis
regulation of membrane potential
urogenital system development
pos. regulation of nervous system development
muscle tissue development
sensory organ morphogenesis
renal system development
pos. regulation of cell projection organization
small GTPase mediated signal transduction
ameboidal−type cell migration
regulation of GTPase activity
regulation of metal ion transport
regulation of developmental growth
muscle system process
1
(4409)
2
(4584)
3
(9419)
4
(8496)
Cluster
p.adjust
1.651686e−22
1.238765e−22
8.258431e−23
4.129216e−23
5.273163e−79
GeneRatio
0.03
0.04
0.05
C
LFC
−4
−2
0
2
4
Ypel2
Akap7
Crebrf
Miga1
Satb1
AW551984
Atxn1
Atpaf1
Bbc3
Atad2b
Rnf44
Afg3l1
Fut8
Plekho1
Meis2
Cxxc5
Gins4
Prr3
Rnf138
Errfi1
Bmper
Acbd5
Myo1b
Elk4
Dact2
Tmem2
Agpat5
Ust
Glis3
Tiam2
Zc3h7b
Cpm
Nol4l
Mycn
Pald1
Kcnj15
St18
Pno1
Smim15
Pfn2
Slc8a1
Dusp4
Kcnk1
Nr4a2
Tenm2
Sox5
Serpinb6a
Nectin3
Foxg1KD
RNAseq
Pantr1KD
ChIPseq
D
aggregated z-score
−4
−2
0
2
4
GATA2,GATA3,GATA1
ZBTB7A,ZBTB7C,PLAGL1,ZBTB7B
RREB1,KLF3,KLF6,KLF4,EGR2, (...)
ZNF524,ZFP524
HIF1A,GMEB2,ARNT,NPAS3,SIM1, (...)
RFX5,RFX6,RFX8,POLR2A,RDBP, (...)
CTCF,RAD21
NR2F1,RXRA,RARG,NR1I2,RARB,HNF4A, (...)
OXG1,FOXK1,FOXE3,FOXB2,FOXJ3, (...)
ZNF143
SPIB,SPI1,ELF1,ETV4,ETV6, (...)
SREBF1
ZNF274,ZFP369,ZFP110
EBF1,SP110,EBF,PRDM16,EBF3, (...)
NFIC,NFIA,NFIX,HIC1,HIC2, (...)
TBX20,TBX10,TBX1,T
GTF3C2,POLR3A,POLR3G
NEUROD6,NEUROD4,NEUROG3,NEUROG2,TCF15, (...)
GM.5.0.Unknown.0017
GATA1,GATA2
FOXI1,FOXE3,FOXB2,FOXK2,ZSCAN16, (...)
TFCP2,TFCP2L1,TCFCP2L1,UBP1,TCFCP2
TCF3,TCF12,ASCL2,MYOD1,TAL1, (...)
1
2
3
4
Cluster

figure S5

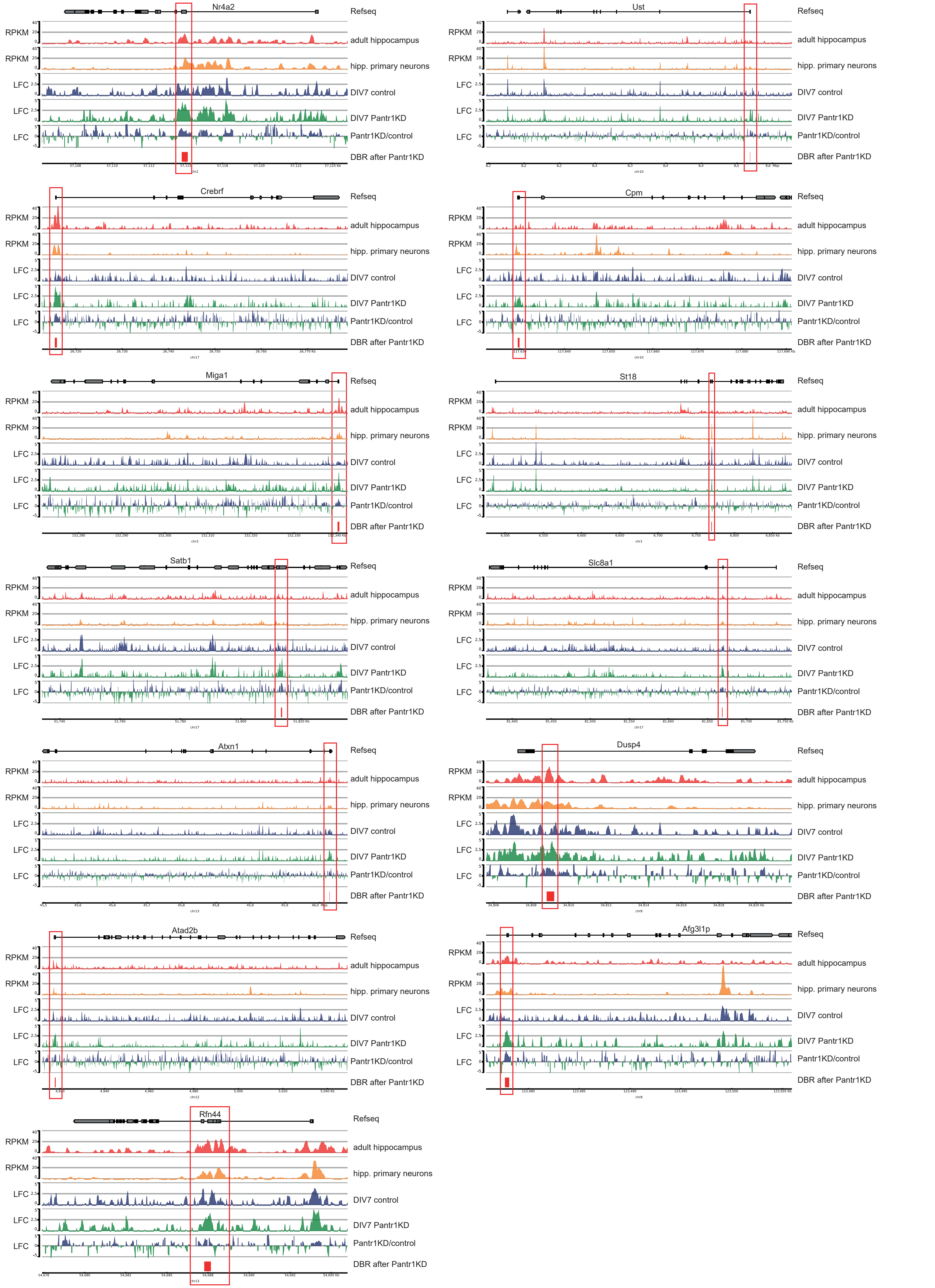
Nr4a2
Ust
Crebrf
Cpm
Miga1
St18
Satb1
Slc8a1
Atxn1
Dusp4
Atad2b
Afg3l1p
Rfn44
Refseq
RPKM
adult hippocampus
RPKM
hipp. primary neurons
LFC
DIV7 control
LFC
DIV7 Pantr1KD
LFC
Pantr1KD/control
DBR after Pantr1KD

figure S6

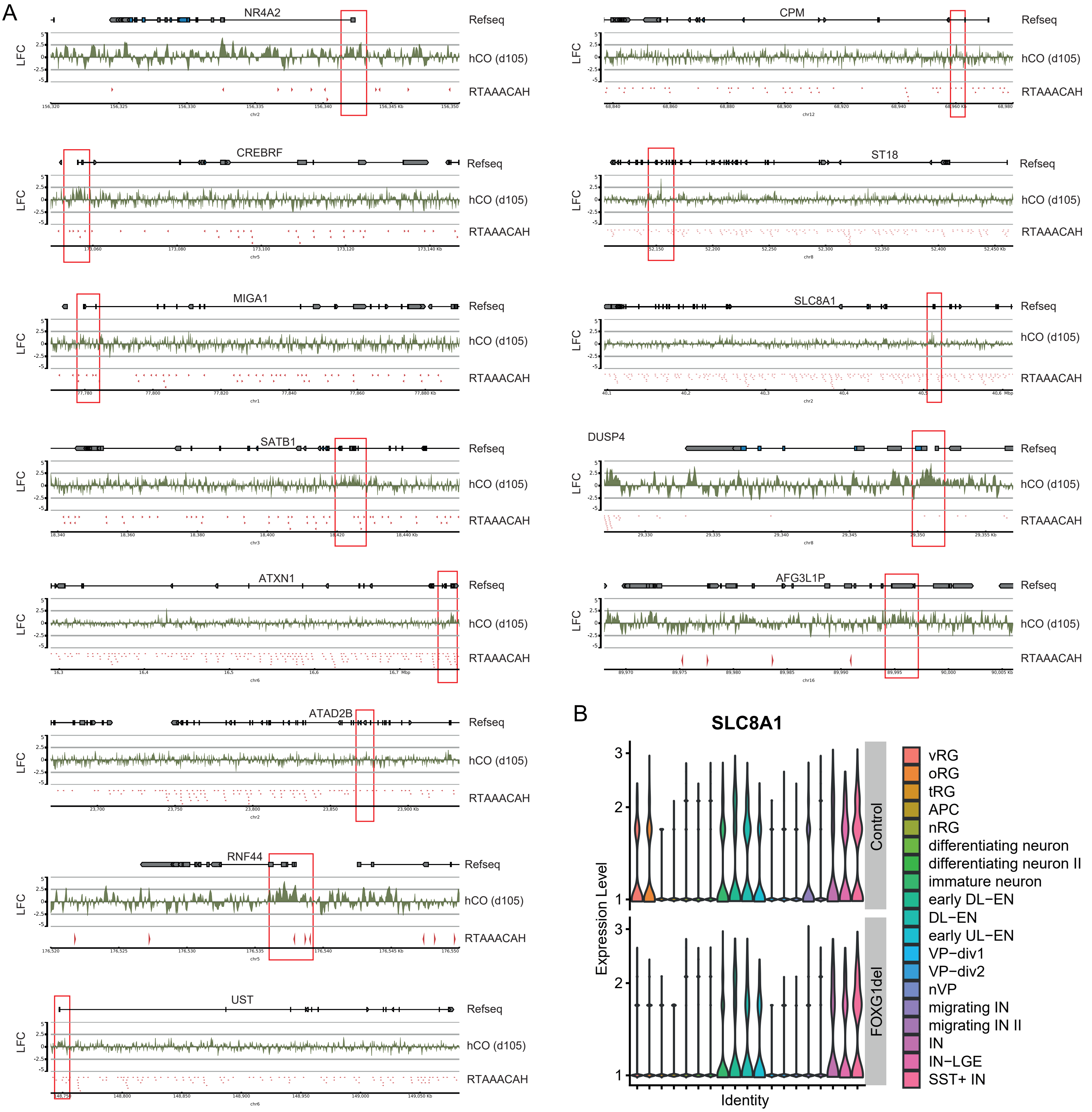
A
NR4A2
CPM
CREBRF
ST18
MIGA1
SLC8A1
SATB1
DUSP4
ATXN1
AFG3L1P
ATAD2B
RNF44
UST
Refseq
LFC
hCO (d105)
RTAAACAH
B
SLC8A1
Expression Level
Identity
Control
FOXG1del
vRG
oRG
tRG
APC
nRG
differentiating neuron
differentiating neuron II
immature neuron
early DL−EN
DL−EN
early UL−EN
VP−div1
VP−div2
nVP
migrating IN
migrating IN II
IN
IN−LGE
SST+ IN

**Supplementary Material and Methods**

*Mouse tissue isolation for RNAseq*

Cortex of E13.5 embryos were dissected from Foxg1$^{cre/cre}$ and wildtype mice (Foxg1$^{+/+}$), hippocampi of E18.5 embryos and adult mice were isolated from Foxg1$^{cre/+}$ and wildtype mice (Foxg1$^{+/+}$). Tissue was subsequently used for RNAseq. Foxg1 lines have C57BL/6J background and are described by the producers [2], (https://www.jax.org/strain/004337).

**Supplemental Table S1*: List of plasmids and their shRNA sequence for lentivirus production***

| **plasmid carrying the coding sequence with short name (bold)** | **sequence for shRNA** | **comment** | **producing company** |
|---|---|---|---|
| pLKO.1-puro-cmv-tGFP sh_Ctrl<br>**shCtrl** | - | used as control for *Foxg1*KD and *Pantr1*KD | MISSION® Luciferase shRNA Control SHC007V MFCD07785395 |
| pLKO.1-puro-cmv-tGFP Sh_Foxg1_1<br>**shFoxg1_1** | CCG GCC TGA CGC TCA ATG GCA TCT ACT CGA GTA GAT GCC ATT GAG CGT CAG GTT TTT G | for *Foxg1*KD | SIGMA MISSION TRCN0000081746 |
| pLKO.1-puro-cmv-tGFP Sh_Foxg1_2<br>**shFoxg1_2** | CCG GCT GAC GCT CAA TGG CAT CTA TCT CGA GAT AGA TGC CAT TGA GCG TCA GTT TTT G | for *Foxg1*KD | Cloned by genescript |
| pLKO.1-puro-cmv tGFP_Sh_Pantr1_1<br>**shPantr1_1** | CCG GGA ATA ACT GCC ATG GAA GGA TCT CGA GAT CCT TCC ATG GCA GTT ATT CTT TTT TG | for *Pantr1*KD | Sigma MISSION TRCN0000179216 |

| | | | |
|---|---|---|---|
| pLKO.1-puro-cmv-tGFP_Sh_Pantr1_2<br>**shPantr1_2** | CCG GGG GAC AGA GTG CCT AGG TAT CTC GAG ATA CCT AGG CAC TCT GTC CCT TTT TG | for *Pantr1*KD | Cloned by genescript |
| pLV(Exp)-EGFP:T2A:Puro-U6>LacZ(shRNA_2)-hPGK>mCherry<br>**Ctrl** | - | control for rescue experiment | Cloned and packaged by Vector builder |
| pLV(Exp)-EGFP:T2A:Puro-U6>FoxG1_shRNA-hPGK>mCherry<br>**shFOXG1_Ctrl** | CCG GCC TGA CGC TCA ATG GCA TCT ACT CGA GTA GAT GCC ATT GAG CGT CAG GTT TTT G | for *Foxg1*KD in rescue experiment | Cloned and packaged by Vector builder |
| pLV(Exp)-EGFP:T2A:Puro-U6>FoxG1_shRNA-hPGK>(Mouse Pantr1_201)<br>**shFOXG1_Pantr1OE** | CCG GCC TGA CGC TCA ATG GCA TCT ACT CGA GTA GAT GCC ATT GAG CGT CAG GTT TTT G | for *Foxg1*KD and *Pantr1*OE in rescue experiment | Cloned and packaged by Vector builder |

**Supplemental Table S2:** ***List of primers for qRT-PCR***

| **target** | **forward primer** | **reverse primer** |
|---|---|---|
| murine Foxg1 | AATGACTTCGCAGACCAGCA | CCGGACAGTCCTGTCGTAAA |
| murine Gapdh | CGGCCGCATCTTCTTGTG | TGACCAGGCGCCCAATAC |
| murine Pantr1 | CGGGACTGTAAGGCGGATAA | GTCCCTCTCCCTCGATGTCA |
| human ACTIN B | CTGGAACGGTGAAGGTGACA | AAGGGACTTCCTGTAACAATGCA |
| human PANTR1 | CATCAGGGGAGCAACGTGAA | TGTCCTGGGAGGCAGTTAGA |
| human FOXG1 | CTGGCGGCTCTTAGAGAT | CCCTCCCATTTCTGTACGTTT |

**Supplemental Table S3:** ***List of antibodies***

| **name** | **usage and dilution** | **company** |
|---|---|---|
| FOXG1 | IHC, 1:500 | Abcam ab196868 |
| FOXG1 | IB, 1:1000, coIP, RIP | active motif #61211 |
| GFAP | IHC, 1:500 | life technologies 13-0300 |
| MAP2 | ICC, 1:200 | Abcam ab32454 |
| NeuN | IHC, 1:150 | abcam ab177487 |
| PAX6 | IHC, 1:150 | BioLegend 901301 |
| TBR1 | IHC, 1:200 | Abcam ab31940 rabbit |

| | | |
|---|---|---|
| TBR2 | IHC, 1:200 | Invitrogen 14-4875-82 |
| TUJ-1 (TUBB3) | IHC, 1:200 | MMS-435P-100 |
| vGLUT1 | IHC, 1:500 | Synaptic Systems (SYSYS) 135 303 |
| vGLUT2 | IHC, 1:500 | Abcam ab79157 |
| anti Mouse-Alexa488 | ICC, IHC 1:500 | Dianova (715-545-151) |
| anti Mouse-Alexa594 | ICC, IHC 1:500 | Dianova (715-585-151) |
| anti Rabbit-Alexa488 | ICC, IHC 1:500 | Dianova (711-545-152) |
| anti Rabbit-Alexa594 | ICC, IHC 1:500 | Dianova (711-585-152) |
| anti Rat-Alexa488 | ICH, 1:500 | ThermoFisher A21208 |
| Streptavidin-Alexa-633 | ICC, 1:500 | Life Technologies S21375 |
| Tidyblot | IB 1:5000 | Biorad Tidyblot |
